# Mapping atherogenesis mechanisms in smooth muscle cells by targeting genes linked to coronary artery disease

**DOI:** 10.1101/2024.11.11.623011

**Authors:** Julián Albarrán-Juárez, Anton Markov, Anne Louise Jensen, Peter Loof Møller, Anna K. Uryga, Djordje Djordjevic, Jakob Hansen, Lise Filt Jensen, Diana Sharysh, Charles Pyke, Jaime Moreno, Giulia Borghetti, Julian Bachmann, Kate Herum, Lisa Maria Røge, Matthew Traylor, Michael Nyberg, Mette Nyegaard, Jacob Fog Bentzon

## Abstract

Recent genome-wide association studies (GWAS) have identified multiple vascular cell-expressed genes linked to coronary artery disease (CAD), suggesting that smooth muscle cells (SMCs) and SMC-derived metaplastic cells are promising targets for novel antiatherosclerosis therapies. However, the disease-promoting pathways of most GWAS-identified genes are unknown, hindering their translation into therapeutic targets. This study integrated public GWAS data for CAD and single-cell RNA sequencing (scRNA-seq) analyses of human atherosclerotic plaques to identify 20 GWAS risk genes with a putative mechanism of action in SMCs or SMC-derived cells.

Gene perturbation experiments in SMCs coaxed to plaque-relevant phenotypes revealed that the selected risk genes, despite encoding very different types of proteins, regulated shared sets of genes associated with contractile functions, cell cycle pathways, NFκB, and type I interferon signaling. By integrating information about GWAS gene effect direction and a deep analysis of cholesterol- and stretch-induced gene modules in SMCs, we find evidence that cholesterol-induced signaling is a pro-atherogenic disease mechanism in SMCs that is upregulated by detrimental and downregulated by protective GWAS genes.

Overall, our study identifies a set of candidate disease mechanisms in SMCs that are regulated by multiple GWAS genes across several SMC assays. Furthermore, it provides proof-of-concept for using GWAS gene effect directionality to predict the pathogenic effect of candidate disease mechanisms that can be extended to other GWAS genes and cell types in the future.

## Introduction

Atherosclerotic cardiovascular disease encompasses myocardial infarction (MI), ischemic stroke, and peripheral artery disease and is the leading cause of death and disability in the world ^1,2^. Atherosclerosis is caused by high levels of low-density lipoproteins (LDLs) and other risk factors (e.g., hypertension and diabetes), which induce inflammation, activation of local arterial smooth muscle cells (SMCs), and the buildup of fibrous, calcified, and necrotic tissue in the arterial intima. After decades of silent development, plaques may suddenly precipitate thrombosis, leading to clinical disease ^3^. Lifestyle changes and pharmacotherapies that halt disease progression by lowering the causal risk factors for atherosclerosis are the central therapies. Despite the availability of these effective interventions, a significant residual risk remains, thus highlighting the need for new approaches that target novel mechanisms.

Vascular SMCs are promising cell targets for new therapies that can work orthogonally to risk factor reduction. Recent murine lineage tracing and human single-cell RNA sequencing (scRNA-seq) studies have shown that local SMCs lose their quiescent contractile phenotype, undergo proliferation and modulate to abundant fibroblast-like and osteochondrocyte-like cells during atherogenesis ^4–8^. Many genes linked to coronary artery disease (CAD) and MI in genome-wide association studies (GWAS) are expressed in SMCs or SMC-derived cells ^9,10^. This indicates that the expansion, modulation, and function of these cells are critical determinants of the outcome of human atherosclerosis ^11–13^. The disease-promoting pathways in SMCs are, however, poorly understood, and this impedes their use as targets for drug development.

In this study, we combined public GWAS for CAD and scRNA-seq plaque data to identify risk genes with a putative mechanism of action in SMCs or SMC-derived cells. By systematically knocking down 20 risk genes that encode very different types of proteins, we performed an unbiased analysis of SMC pathways that are genetically linked to human atherosclerosis, and therefore, they represent potential targets for SMC-directed therapies.

## Results

### Identification of GWAS genes with a potential mechanism of action in SMCs

A crude list of 242 genomic variants associated with CAD independently of LDL cholesterol was drawn from the Cardiovascular Disease Knowledge Portal (CVDKP) and FinnGen study (Freeze 5) (**Figure 1A** and **Supplementary Table 1).** Combined data from CVDKP, FinnGen, and Open Targets Genetics platform suggested 368 genes to be putatively regulated by the collected variants. To identify candidate GWAS genes with a potential function in plaque SMCs or SMC-derived cells, we looked up the expression of these genes in human plaque scRNA-seq data, assembled by integrating published data sets of human coronary ^5^ and carotid ^6,7^ atherosclerosis (**Figure 1B**). The integrated dataset comprised 50,390 cells after quality control and filtering, including a large mesenchymal supercluster (15,018 cells) of cells with SMC, fibroblast, pericyte, transitional, fibromyocyte, and osteochondrogenic phenotype based on marker gene expression (**Figure 1C** and **Supplementary Table 2).** Of the 368 examined genes, 92 protein-coding genes showed enriched expression in the mesenchymal supercluster compared with other cell types (**Figure 1D and Supplementary Table 1)**. From this pool, we selected 17 genes (*ADAMTS7, C1S, COL4A2, CRISPLD2, CTTN, DSTN, FHL1, GEM, HHIPL1, LMOD1, LOXL1, LRP1, MFGE8, PDGFD, PLPP3, TIMP2*, and *TNS1*), chosen to represent a wide range of putative functions **(highlighted in Figure 1D**). The list included genes with strong expression in mesenchymal subclusters (e.g., *C1S, DSTN, MFGE8*), genes with experimental evidence of functions in SMCs (e.g., *LMOD1, PDGFD*)^14–16^, and genes encoding both extracellular (*PDGFD, C1S, COL4A2, HHIPL1, LOXL1, ADAMTS7, MFGE8, CRISPLD2, TIMP2*) and intracellular (*DSTN, CTTN, FHL1, ALDH2, LMOD1, GEM, RBPMS2, PLPP3, TNS1, LRP1)* proteins, according to the human protein atlas ^17^. We added three CAD GWAS genes (*ALDH2, TCF21, RBPMS2*) with published evidence for a function in SMCs ^5,18,19^. Lead genomic variants and plaque single-cell expression profiles for the 20 selected genes are shown in **Supplementary Table 3** and **Supplementary Figure 1**, respectively.

**Figure 1:**
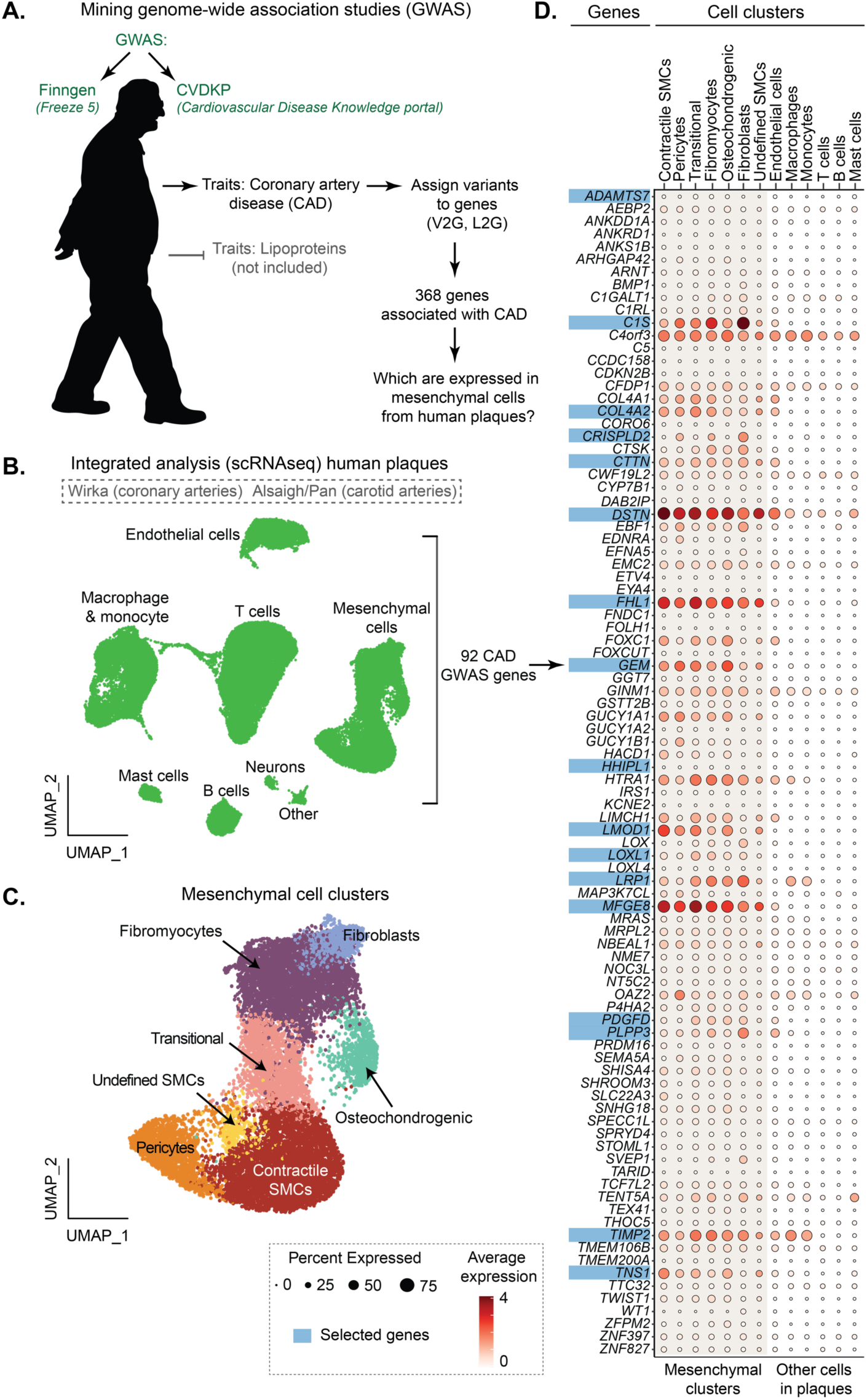
From GWAS and scRNA-seq data to the selection of genes. **A.** GWAS results from Finngen and CVDKP were utilized to identify variants associated with coronary artery disease (CAD), excluding those associated with lipid traits. After gene fine-mapping, we identified 368 genes. **B.** We then tested the expression of these genes on integrated single-cell transcriptomic data from publicly available studies of human atherosclerotic lesions (in total, 50390 cells from 10 patients). This led to identifying 92 protein-coding genes preferentially expressed in mesenchymal clusters (25456 cells). **C**. Shows a supercluster of mesenchymal cells was subdivided into 7 clusters that were identified by their marker genes as contractile smooth muscle cells (SMCs), transitional, osteochondrogenic, and undefined SMCs, pericytes, fibromyocytes, and fibroblasts in integrated data. **D.** Dot plot shows the expression of genes enriched in the mesenchymal supercluster. Normalized and log2-transformed gene expression values are averaged for every cell cluster.

Expression of the selected target genes was also investigated in a spatial transcriptomics data set of human coronary plaques ^20,21^. *MFGE8, TIMP2, TNS1, C1S, COL4A2, DSTN, LMOD1,* and *LRP1* were abundantly expressed in coronary artery plaque and underlying media. *ALDH2, CTTN, GEM, LOXL1, PLPP3*, and *RBPMS2* expression were restricted mainly to the media, while *CRISPLD2, TCF21, HHIPL1,* and *ADAMTS7* did not have sufficient detectable expression to determine the localization (**Supplementary Figure 2**).

### Establishment of cellular assays to evaluate human smooth muscle cell function

To test gene function in several SMC phenotypes that are relevant in atherosclerotic plaques, we coaxed human aortic SMCs into three different cellular assays: baseline, cholesterol, and mechanical stretch (**Figure 2A**). The underlying rationale was that target genes that might drive important functions in one context may be inactive in another. The baseline condition was a standard SMC culture, where cells spontaneously modulate, losing contractile proteins, increasing proliferation, and displaying a less elongated morphology ^22,23^. Secondly, SMCs were overloaded with cholesterol to drive cholesteryl ester droplet formation in SMCs ^24^, which leads to a foam cell-like state characterized by increased inflammatory signaling ^25,26^. Furthermore, mechanical stretching (10% elongation) was applied to mimic the physiological mechanical forces of the arterial wall and potentially drive SMCs toward a less inflammatory and more contractile phenotype ^27^. RNA-seq analysis confirmed that 72h of cholesterol overloading in SMCs upregulated inflammatory genes (e.g., *CCL20, CXCL2, CXCL3, CXCL5, IL1B*), cholesterol efflux genes (*ABCA1* and *ABCG1*) and downregulated contractile genes (e.g., *ACTA2, CNN1, LMOD1, MYOCD*, and *TAGLN*) compared with cells cultured under baseline conditions (**Figure 2B**). In contrast, 6h of physiological mechanical stretching downregulated several inflammatory genes (e.g., *CCL2, CXCL1, CXCL3, IL1B, IL6*) and upregulated contractile genes (e.g., *CNN1, ACTA2, TAGLN, ACTG2*) compared with SMCs cultured similarly under static conditions (**Figure 2C**).

**Figure 2:**
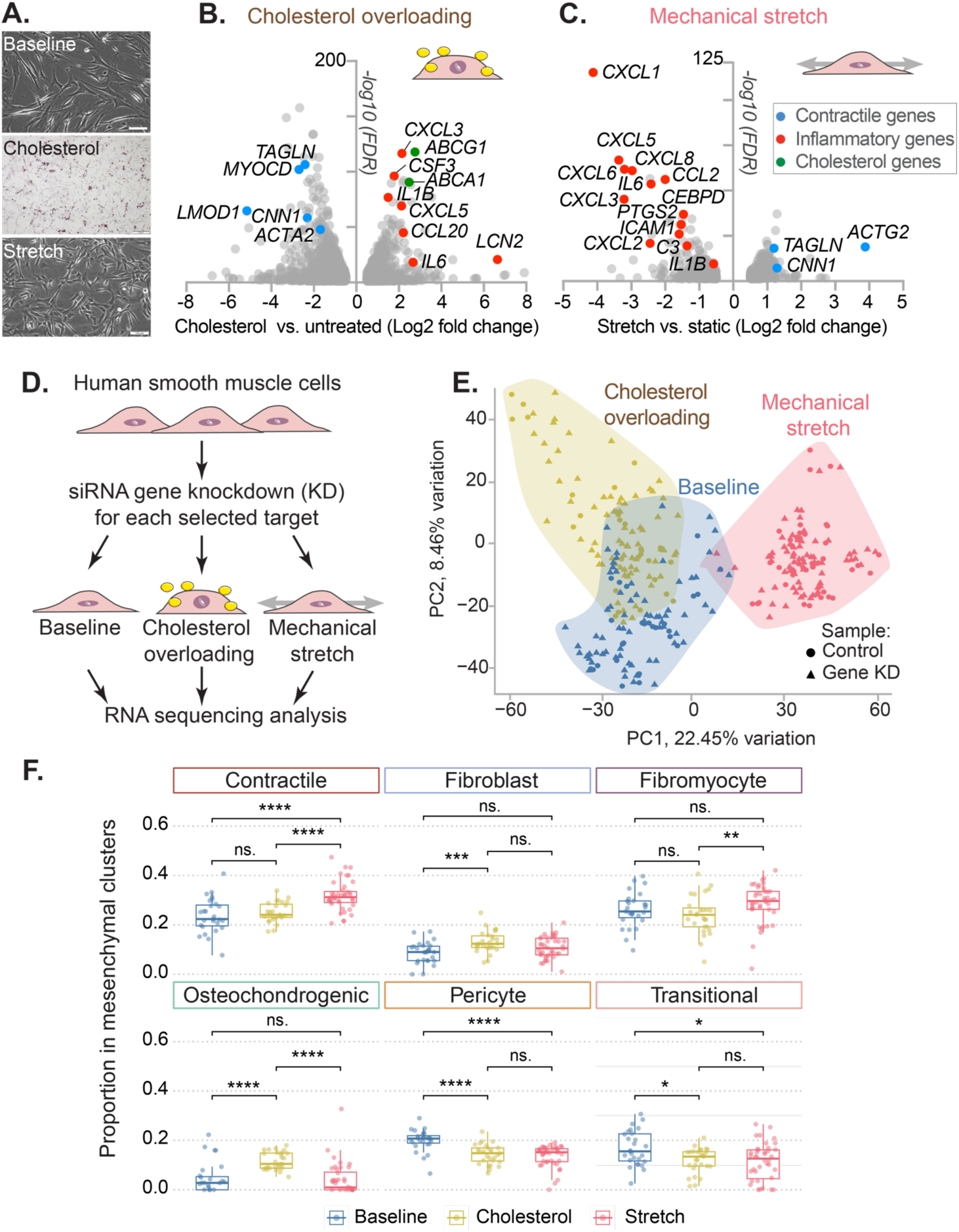
*In vitro* models of vascular disease in human smooth muscle cells. **A.** Representative brightfield images of human smooth muscle cells at baseline or after cholesterol overloading or 10% mechanical stretch. Scale bars 100 um (baseline and stretch), 200 um (cholesterol). **B.** Gene expression in human smooth muscle cells after cholesterol overload (volcano plot). **C.** Gene expression in SMCs after physiological (10%, 1hz, for 6 h) mechanical stretch (volcano plot). A and B show significantly regulated genes (*P value* < 0.05 and absolute log2 fold change ≥ 0.5) compared to their control condition (3 technical replicates in each group). FDR. False discovery rate-adjusted values. **D.** Overview of the workflow. SMCs were transfected with siRNAs for each gene, subjected to different cellular assays, and analyzed by RNA sequencing. **E.** Principal component analysis (PCA) of transcriptome profiles of studied SMC samples (*n*=345) revealed the most difference between mechanical stretch and other cellular assays. **F.** Bulk RNA-seq data from SMCs transfected with control siRNA during baseline (*n*=27), cholesterol overload (*n*=27), and stretch (*n*=40) conditions were decomposed into components corresponding to mesenchymal cell types annotated in single-cell RNA-seq data of human atherosclerotic plaques. The undefined cluster was excluded from this analysis. P values are shown according to the Wilcoxon-Mann-Whitney test and adjustment for multiple comparisons. * Adjusted P value < 0.05, ** P value < 0.01, *** P value < 0.001, **** P value < 0.0001 or ns. (not significant).

### Knockdown of genes associated with coronary artery disease in human SMCs

To analyze the function of the selected CAD risk genes in human SMCs, we knocked down each target gene by custom-designed small interfering RNAs (siRNAs) and subjected cells to baseline, cholesterol overload, or mechanical stretch conditions. Effects on gene expression were analyzed by RNA sequencing using a nested design where 2-3 target gene knockdowns were run concurrently with control siRNA-treated cells to mitigate batch effects (**Figure 2D**). Overall, we analyzed the transcriptome of 345 samples (3 to 4 technical replicates per target gene and condition). The numbers of differentially expressed genes (DEGs) compared with controls were higher under baseline conditions (from 159 to 3266 DEGs, 884 at average) and cholesterol loading (from 107 to 3122 DEGs, 1078 at average) than after mechanical stretch (from 12 to 1612 DEGs, 592 at average). Principal component analysis (PCA) of total RNA sequencing results confirmed the transcriptional changes across the baseline, cholesterol overloading, and mechanical stretch conditions (**Figure 2E**). It also revealed considerable overlap between baseline and cholesterol loading conditions. To predict which mesenchymal cell states observed *in vivo* in atherosclerotic lesions can be mirrored by *in vitro* samples in our experiments, we performed cell-type deconvolution analysis of bulk gene expression data from SMCs across our cellular assays. First, we generated pseudo-bulk samples based on scRNA-seq data from human atherosclerotic lesions and verified that the proportions of mesenchymal cell clusters in these pseudo-bulk samples, as estimated by the cell type deconvolution method Bisque, were similar to those observed in the original scRNA-seq dataset (**Supplementary Figure 3**). Then, we estimated mesenchymal cell proportions for SMC samples transfected with control siRNA and compared them across our cellular assays. This analysis revealed that SMCs subjected to stretch show a higher proportion of contractile components than baseline and cholesterol. Similarly, cells under cholesterol overloading increased their content of osteochondrogenic components compared to baseline or stretch. In addition, cells under baseline conditions showed increased proportions of transitional SMC components compared to cholesterol or stretch (**Figure 2F**). This analysis confirms that our cellular assays recreate, to some degree, different cell states observed *in vivo* in atherosclerotic lesions.

The knockdown efficiency of the target genes was analyzed by real-time qPCR (qPCR), as shown in **Supplementary Figure 4.** Gene knockdown analysis was efficient for most target genes, decreasing the level of target gene expression by 75 to 90% compared to control samples. However, for *TCF21* and *PLPP3,* the designed siRNAs reduced the expression by less than 50%. Therefore, *TCF21* and *PLPP3* were not further analyzed in the study. RNA-seq data confirmed the knockdown efficiency and interestingly showed mutual regulation among the target genes (**Supplementary Figure 5**). For example, the knockdown of *LMOD1* significantly upregulated the expression of *GEM, CTTN, and TNS1* and downregulated the expression of *ALDH2, COL4A2, CRISPLD2, FHL1, PDGFD, and TIMP2* in baseline and cholesterol overloading conditions.

### Regulation of smooth muscle cell pathways after target gene knockdown

Gene set enrichment analysis (GSEA) for all comparisons in this study showed that cell cycle, interferon signaling, inflammatory cytokines, and Rho GTPases were among the top gene ontology categories and biological pathways regulated after target gene knockdowns (**Supplementary Table 4)**. For a more direct analysis of pathways and genes relevant to SMC functions, we analyzed tailored gene signatures representing SMC contraction, cytoskeleton, cell adhesion, extracellular matrix organization, cell cycle, and inflammatory/immune responses, collected from several sources, including KEGG ^28^, Reactome ^29^, and WikiPathways ^30^ **(Supplementary Table 5).** Gene sets involved in SMC contraction, extracellular matrix synthesis, and cell cycle were regulated by multiple target genes, and we also identified a strong regulation of the nuclear factor kappa-light-chain-enhancer of activated B cells (NFκB) and interferon pathways (**Figure 3A-C**).

**Figure 3:**
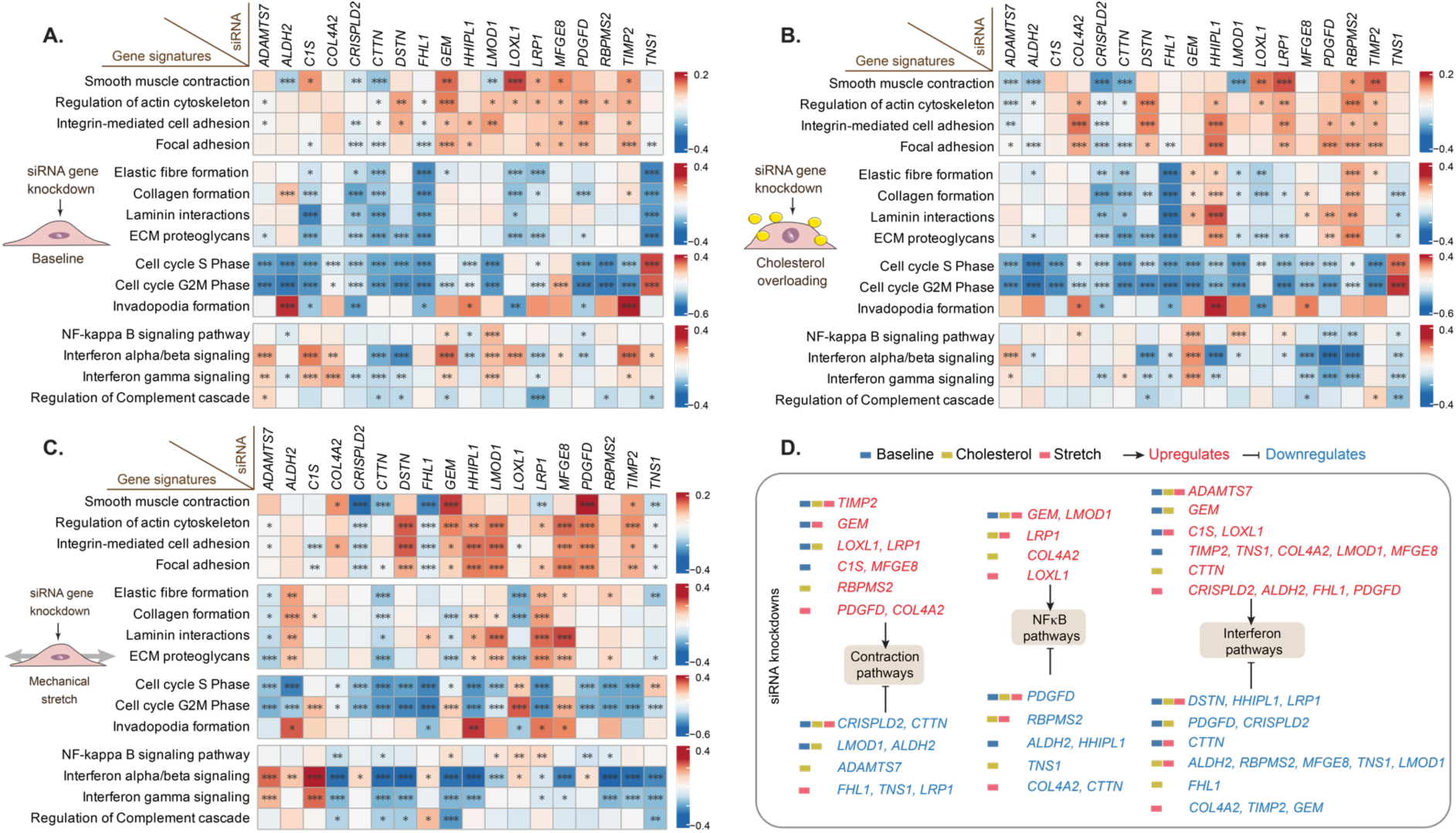
Gene signatures disturbed after target gene knockdown. Expression-based scoring of gene signatures implicated in SMC contractility, synthesis of extracellular matrix proteins, cell cycle, and inflammatory response in the different cellular assays: **A.** baseline, **B.** cholesterol overload, and **C.** mechanical stretch. The color scale represents a z-score of deviation in the transcriptional activity of relevant pathway genes in target knockdown compared with a control group (3 or 4 technical replicates in each group). Asterisks highlight statistical significance based on false discovery rate (FDR)-adjusted P values (Padj): * Padj <0.1, ** Padj <0.01; *** Padj <0.001. Extracellular matrix (ECM) **D.** Examples of pathways regulated by differentially expressed genes induced by siRNA knockdowns of target genes.

For a subset of target genes (e.g., *ADAMTS7, CTTN, GEM, LMOD1, PDGFD, TIMP2*), the regulation of specific signaling pathways remained consistent across all examined cellular states, but often, it was only detected in one or two of the examined cell states and, in some cases (e.g., *LRP1, COL4A2, and TNS1*), in opposite directions, as exemplified for contraction, NFκB, and interferon signaling in **Figure 3D**. This may indicate that certain mechanisms gain or wane in importance in different cell states. For example, genes involved in extracellular matrix organization were generally downregulated by knockdowns in the baseline conditions, presumably reflecting the high activity of these genes in this cell state, while up- and downregulation were seen in other states (**Figure 3A-C**). Basal expression of the target genes, although in several cases up- or downregulated by cholesterol or stretch (**Supplementary Figure 6**), did not predict knockdown outcomes.

To validate the observed regulation of NFκB- and type I interferon-induced genes, we performed two types of experiments. First, we tested whether the siRNA transfections themselves could cause the observed inflammatory signaling. Depending on specific recognition motifs in the sequence, some siRNAs can directly activate type I interferon responses ^31,32^. The siRNAs used in our experiments were designed not to contain these motifs, and we confirmed experimentally that neither the control siRNA nor lipofectamine transfection alone altered the expression of *IL6, IFI6, IFITM1,* or the contractile marker *MYOCD* compared with untreated controls **(Supplementary Figure 7)**.

Second, since genes induced by NFκB and type I interferon signaling may vary by cell type, we defined the NFκB and type I interferon gene signatures in human SMCs by stimulating with the prototypical inducers tumor necrosis factor (TNF) and interferon alpha (IFN-alpha) followed by RNA sequencing **(Supplementary Figure 8A-B** and **Supplementary Table 6)**. We found that the experimentally defined gene sets were partly overlapping and confirmed that they were regulated by multiple target genes with *C1S* and *HHIPL1* knockdown, producing the strongest up- and down-regulation across the three cell states **(Supplementary Figure 8C)**.

### Widespread alterations in inflammation- and cell cycle-related regulons

To identify the transcription factors (TFs) that drive the shared transcriptomic changes among different knockdowns, we detected regulons (a set of genes jointly regulated by the same transcription factor) among up and downregulated genes. To focus on regulons with *in vivo* relevance, we restricted to regulons that were detected to be active in at least one mesenchymal cell subcluster of human plaques by scRNA-seq analysis with SCENIC (**Figure 4A**). Corroborating our analysis on the pathway and marker gene levels, we found that regulons putatively controlled by cell cycle/proliferation (E2F3, E2F4, ETS1, NFATC1, SP1, MEIS1, and TAF1) and NFκB/interferon (IRFs, STAT, RELA, RELB, NFKB1) transcription factors were significantly enriched. Members of the AP-1 complex (FOS, JUN), SOX, and Kruppel-like factors (KLFs) known as regulators of SMC phenotype were also identified (**Figure 4B**). A detailed list of the regulons identified in every targeted gene knockdown and cellular assay is shown in **Supplementary Table 7.**

**Figure 4:**
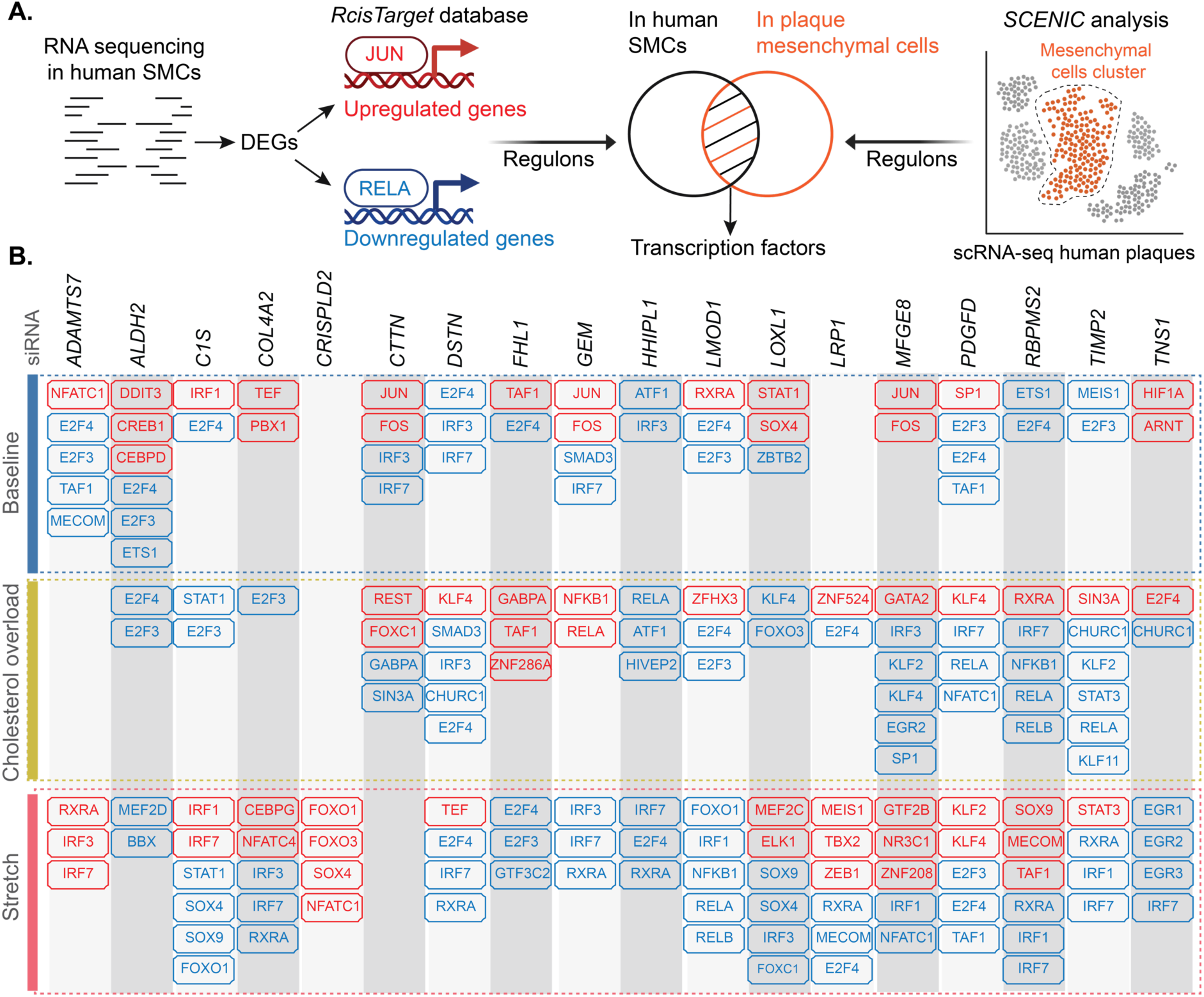
Regulons with altered activity following target knockdown in different cellular assays. **A.** Regulon is a collection of genes with a shared transcription factor binding site suggesting potential co-regulatory mechanisms by identical transcription factors (TFs). For each targeted gene knockdown and corresponding cellular assay in human SMCs, we identified regulons with a significant presence within the differentially expressed genes (DEGs), separately for up- and downregulated gene sets (JUN and RELA are shown as examples). Identified regulons were then overlapped with regulons detected to be active in at least one mesenchymal subcluster by SCENIC analysis of the integrated single-cell RNA sequencing dataset from human plaques. Drawing created with Biorender.com. **B.** An overview of shared regulons across various mesenchymal cell clusters and altered by targeted gene knockdown in cellular assays are named by putatively binding TFs. Regulons with up-and down-regulated genes are depicted as red and blue boxes, respectively.

### Independent regulation of common SMC marker genes and functional readouts

The growth of SMC-derived cells in atherosclerosis involves loss of SMC contractile gene expression, increased proliferation, and switching to modulated cell phenotypes characterized by extracellular matrix production and inflammatory signaling ^8,33,34^. This metaplastic process explains the inverse correlation between contractile genes and markers of modulated cells and inflammatory signaling in plaque scRNA-seq data ^8^. It is, however, less clear if reciprocal regulation of these programs already exists within a particular SMC cell state. To analyze this, we retrieved the expression levels of multiple marker genes used in the literature to define SMC phenotype, including contractile gene markers (*ACTA2, ACTG2, CNN1, LMOD1, MYOCD, TAGLN*), modulated SMC markers (*FN1, KLF4, LUM, SPP1*), cell proliferation markers (*MKI67, PCNA*), NFκB activation markers (*IL1B, IL6, PTGS2, VCAM1*) and type I interferon-induced genes (*IFIT1, IFITM1, ISG15, MX1, OAS1*).

Contractile marker gene expression was widely regulated under baseline conditions, being stimulated by *C1S, GEM, LOXL1, MFGE8, RBPMS2,* and *TIMP2* knockdown and repressed by *ADAMTS7, ALDH2, COL4A2, CRISPLD2, CTTN, DSTN, FHL1, HHIPL1, LMOD1,* and *PDGFD* knockdown (**Figure 5A**). Changes in cholesterol-overloaded and stretched cells were similar albeit with some differences in the magnitude of regulation. The selected contractile genes were mostly regulated as a group, consistent with their common transcriptional regulation ^35^, with *MYOCD* showing the most independent regulation.

**Figure 5:**
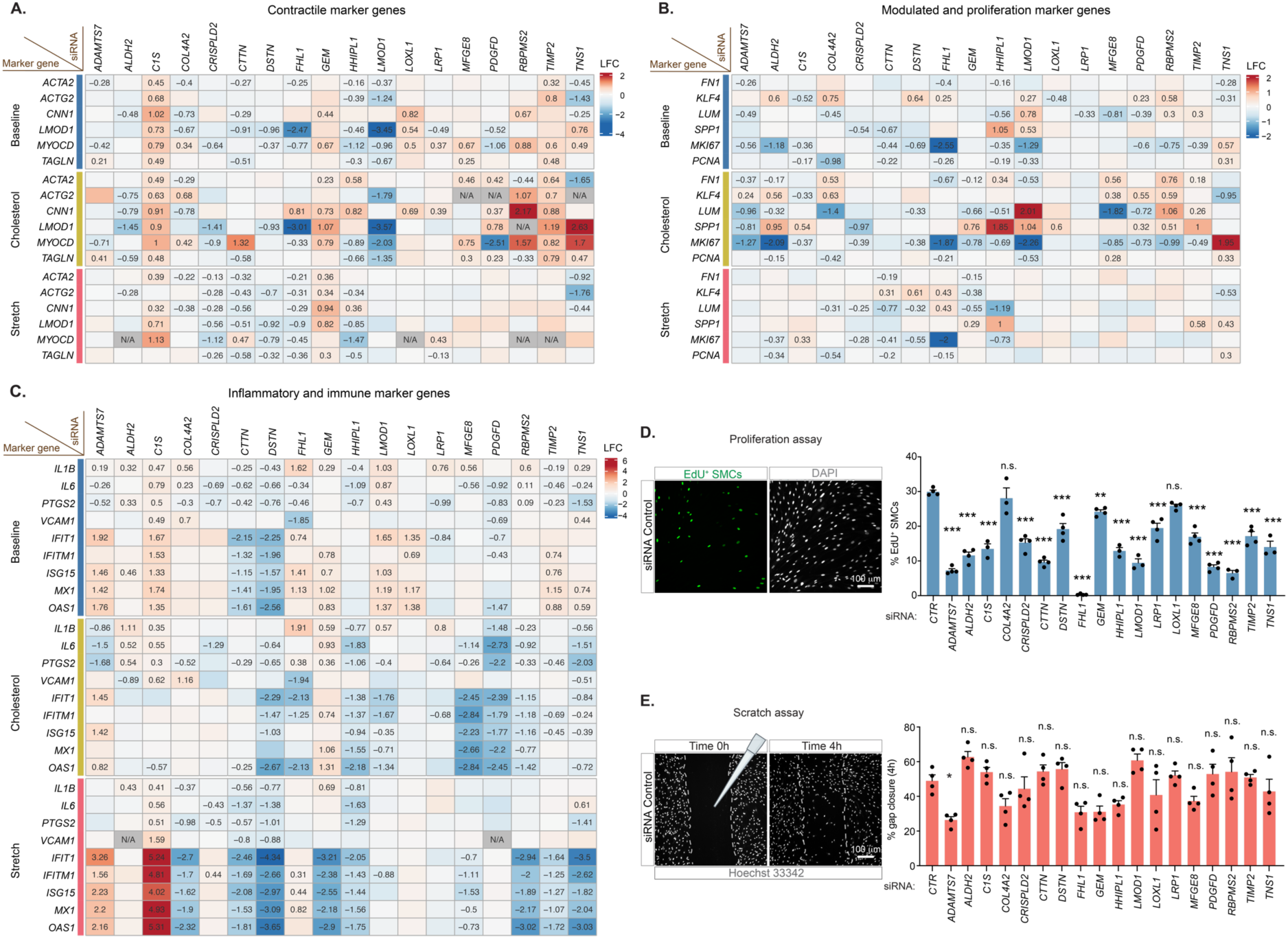
Regulation of conventional gene markers of smooth muscle cell function. Differences in the expression of a subset of marker genes important for smooth muscle cell function in target gene knockdowns compared to related control groups on each cellular assay. Results are shown for different cellular pathways: **A.** contractile, **B.** modulated and proliferation, and **C.** inflammatory and immune marker genes. The number in heatmaps indicates log2-transformed fold changes (LFC). Only statistically significant LFC numbers are shown. N/A indicates that gene expression was not detected. **D.** SMCs were transfected with negative control siRNA (control) or siRNAs targeting each gene and labeled with EdU (5-ethynyl-2’-deoxyuridine) for 24 h to evaluate their proliferation. Shown is a representative image of the technique and the percentage of EdU+ SMCs per view field (3-4 technical replicates were evaluated per gene). **E.** SMCs were transfected with the different siRNAs, and a scratch was made in the center of the well (representative image is shown). Cell nuclei were labeled with Hoechst 33342 live staining (NucBlue^TM^) and incubated at 37°C and 5% CO2 in a time-lapse microscope (Nikon, Eclipse Ti2). Cell migration was monitored for 24 hours in total. The percentage of gap area closure is shown 4 h after the scratch (3 to 4 technical replicates were evaluated per gene). Samples were compared to the control cell using one-way ANOVA followed by a Dunnett test. A * represents an adjusted P value < 0.05, ** P value < 0.1; and *** P value < 0.01, or n.s. (not significant, P value > 0.05) compared to control.

Interestingly, the selected markers for modulated SMCs (*FN1, KLF4, LUM*) were often independently regulated and not necessarily upregulated when contractile genes were repressed and vice versa. (**Figure 5B**). Neither did we observe a consistent reciprocal regulation between contractile genes and marker genes for proliferation (*MKI67, PCNA)*, which were downregulated after all target gene knockdowns except for *TNS1* (**Figure 5B**). Moreover, while markers for NFκB signaling and type 1 interferon signaling were regulated by multiple target genes (**Figure 5C**), those knockdowns that dampened inflammatory signaling, e.g., *DSTN* and *CTTN*, were not necessarily the same that upregulated modulated SMC markers and vice versa.

Finally, we examined proliferation and migration, which are functional readouts often used to gauge SMC phenotype. Proliferation measured by EdU labeling was downregulated in almost all target gene knockdowns (except for *COL4A2* and *LOXL1*, which showed no difference), consistent with the cell cycle marker gene analysis (**Figure 5D**). In contrast, a significant difference in cell migration was only detected for *ADAMTS7* knockdown (**Figure 5E**).

Overall, this analysis, made possible by having a large set of independent cell perturbations, indicated that contractile gene expression and markers/functional readouts of proliferation, modulation, and inflammatory signaling are not tightly co-dependent in cultured SMCs. This is important because such markers are often used as markers for SMC phenotypic switching in cell culture under the assumption that they are linked and report cell identity changes similar to those occurring during atherosclerosis, which appears not to be justified.

### Proatherogenic effect of cholesterol overload indicated by GWAS gene effect directions

The genetic variation identified in GWAS studies mostly influences CAD risk by altering gene expression rather than through alterations in the encoded protein, and genes can be detrimental or protective depending on whether lower expression decreases or increases disease risk, respectively. To classify our selected target genes, we identified co-localizing expression or protein quantitative trait loci (e/pQTLs) and determined their impact on CAD risk by Mendelian randomization ^36^. We used eQTL data from GTEx V8 ^37^ and eQTLgen ^38^, and pQTL data from UK Biobank ^39^ and in a large Icelandic cohort ^40^.

The analysis predicted *ALDH2, C1S, CTTN, LOXL1, MFGE8, PDGFD, and TIMP2* to be detrimental and *FHL1, HHIPL1, LMOD1, LRP1, RBPMS2* to be protective genes for CAD development **(Supplementary Table 8)**. The analysis was inconclusive for the residual genes. We then used this classification as a tool to estimate which of the many regulated pathways identified in our SMC analysis were likely pro-versus anti-atherogenic. The underlying rationale is that a pro-atherogenic mechanism will tend to be upregulated by the knockdown of protective genes, while an anti-atherogenic mechanism will tend to be upregulated by the knockdown of detrimental genes.

Analyzing the broader gene signatures induced by cholesterol overloading and stretch revealed that cholesterol overloading in SMCs had the characteristics of a pro-inflammatory mechanism (**Figure 6A**): knockdown of protective genes, such as *LMOD1, FHL1, or LRP1,* upregulated the cholesterol overload-induced gene signature, while knockdown of detrimental genes, such as *C1S* and *TIMP2*, downregulated it. Interestingly, this pattern of opposite regulation by protective and detrimental genes was not seen among gene knockdowns that simultaneously upregulated stretch-induced genes, suggesting some level of interaction (e.g., *HHIPL1, MFGE8*).

**Figure 6.**
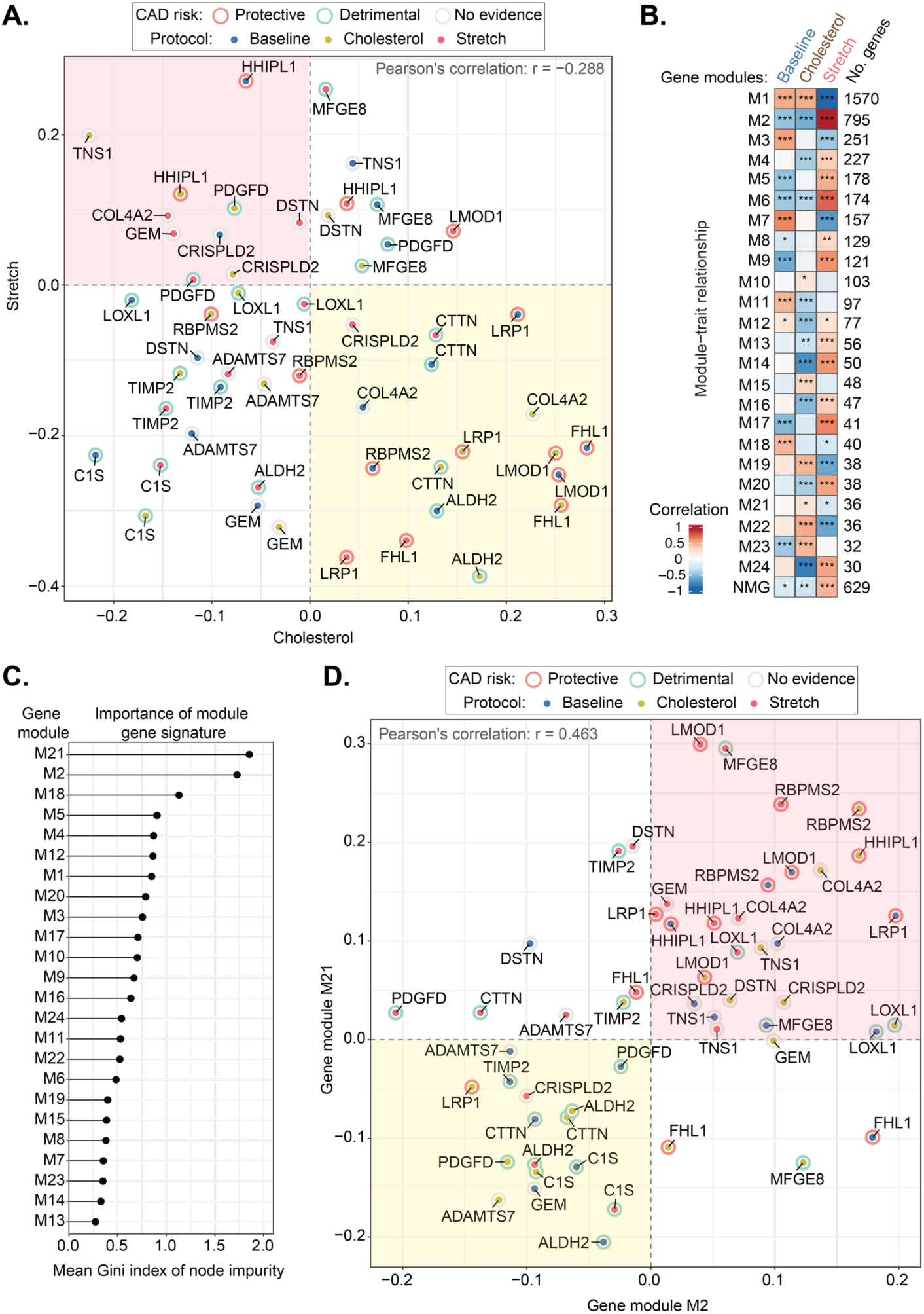
Gene signatures in human SMCs. **A.** Regulation of cholesterol and stretch-specific gene signatures (top 50 uniquely up-regulated genes after either cholesterol overload or stretch) scored in target gene knockdowns. The type of cellular assay (baseline, cholesterol overload, or stretch) is marked by dots and the estimated effect of the target gene on human CAD (protective, detrimental, or unknown) by circles. **B.** Correlation of co-expressed gene modules with the cellular assays. The color scale on the heatmap represents the Pearson’s correlation coefficient between module eigengene (summarized expression profile) and coaxed cellular state. Asterisks highlight significant correlation: * P value <0.05, ** P value <0.01; *** P value <0.001. NMG, Non-module genes not clustered in any co-expressed module. **C.** Random Forest model and Gini index-based estimate of feature importance were used to prioritize identified module gene signature scores by their ability to discriminate between target genes with detrimental and protective effects on CAD. The higher the estimate, the better the gene module score predicts the opposite effects of target genes. **D.** Relationship between the top gene modules (M02 and M21) gene signature dynamics in target gene knockdowns, genetic effect direction of target genes, and cellular assay. The top 100 highly connected genes used in the analysis for the large M2 module.

Individual pathways (e.g., SMC contractility, NFκB signaling, type I interferon) were not consistently regulated by protective versus detrimental genes (analyses not shown). This may reflect the complex and mutually independent regulation of these pathways across the gene knockdowns, necessitating a multivariate analysis to disentangle that was not possible with our limited number of informative genes.

### Specific SMC gene modules regulated by protective and detrimental GWAS genes

To evaluate which parts of the cholesterol- and stretch-induced responses may be pro- or anti-atherogenic, we defined gene modules in SMCs subjected to baseline, cholesterol overload, and mechanical stretch conditions by analyzing gene co-expression in the 87 samples transfected with control siRNA. We obtained 24 modules (M1 to M24), ranging in size from 1570 genes in M1 to 30 genes in M24, with evidence of differential regulation across the SMC states (**Figure 6B**). Details of genes and their connectivity are provided in **Supplementary Table 9-10,** and a graphical representation of the analysis can be found in **Supplementary Figure 9.**

We then explored which of these 24 gene modules meets the criteria of a pro- or anti-atherogenic mechanism, being regulated oppositely (if at all) by protective and detrimental genes. Gene module 2 (M2) and module 21 (M21) were the top modules identified with these characteristics (**Figure 6C**). Module 2 (795 genes) was characterized by genes with high activity in the stretch condition but low in baseline and cholesterol-overloading conditions. In comparison, module 21 (36 genes) contained genes with high activity during cholesterol-overloading conditions but not at baseline or stretch. Enrichment analysis revealed that module 2 contained genes involved in RNA processing, while module 21 comprised genes associated with the wingless-type MMTV integration site family (Wnt) signaling pathway (**Supplementary Table 10**). Expression of the two modules across the target knockdowns showed a positive correlation, and both had the signature of a pro-atherogenic mechanism being upregulated by protective gene knockdowns (e.g., *LMOD1, HHIPL1*) and downregulated by detrimental gene knockdowns (e.g., *ALDH2, C1S*) (**Figure 6D**). These elements of SMC responses to metabolic and mechanical stress may, therefore, be particularly interesting for further exploration. An expression-based gene scoring analysis for all the modules is shown in **Supplementary Figure 10.**

## Discussion

Many CAD susceptibility genes in GWAS studies do not exert their effects through classical risk factors such as circulating cholesterol levels or hypertension ^41^. Therefore, unidentified disease mechanisms contributing to residual risk, which are not addressed by current therapies, may exist. During atherosclerosis, a subset of SMCs de-differentiate, proliferate, and modulate to other mesenchymal cell phenotypes, many of which abundantly accumulate cholesteryl ester lipid droplets ^8,42,43^. The accumulation of the SMC-derived cells and the extracellular matrix proteins they secrete is a central mechanism by which atherosclerotic plaque grows and a main determinant of plaque stability ^44^. Cells with SMC-like phenotype build the fibrous cap, which protects against plaque rupture and thrombosis, but the modulated SMCs in the plaque interior may play detrimental roles by contributing to the formation and expansion of the necrotic core ^45,46^. The phenotypic transition of SMCs and the important functions of the modulated cell types in plaques make them interesting therapeutic targets, but little is known about the molecular pathways that could be targeted.

Recent studies have searched for genes regulating a fixed set of atherosclerosis-relevant SMC functions (e.g., proliferation, inflammatory gene expression) using the genetic variation across a large set of patient-cultured SMCs ^47,48^. Here, we followed the complementary strategy of introducing gene perturbations experimentally and performing an open search for alterations in SMC gene expression and function. This has previously been done for individual genes, but as exemplified by our findings, many patterns of atherosclerosis-relevant SMC regulation can only be revealed by looking at multiple gene perturbations in the same context. We systematically performed a knockdown screen of genes that are expressed in SMCs or their modulated progeny in coronary and carotid plaque scRNA-seq datasets and have genetic evidence for a causal role in CAD that is not explained by plasma lipids. The goal was to use these knockdowns as an instrument to understand what gene signatures and pathways are relevant for the pathophysiological function of SMCs in CAD. While it is not yet possible to recreate the SMC phenotypic transitioning process in vitro, we steered human SMCs to several plaque-relevant states to increase the likelihood that central pro- or anti-atherogenic mechanisms of SMCs were featured in our assays. Finally, by integrating what is known about the effect on CAD of gene variants controlling the expression of our GWAS target genes, we attempted to determine which of the observed regulated pathways are pro-atherogenic and anti-atherogenic.

Most of the genes targeted in our study have not previously been investigated for their role in human SMC function or modulation (e.g., *DSTN, CTTN, FHL1, GEM, TNS1*, etc.), but some have been examined in independent studies, which can be used for cross-validation of our data ^49–55^. PDGFD was shown previously to promote the modulation of cultured SMCs and increase inflammatory signaling ^14^, while LMOD1 was required for the maintenance of the contractile phenotype ^15,16^. These findings were replicated in our data. The knockdown of *PDGFD* decreased the expression of genes related to NFκB/interferon signaling and collagen synthesis, while the knockdown of *LMOD1* reduced genes related to SMC contractile gene expression.

Overall, our analysis produced four key findings. Firstly, we observed widespread regulation of cytoskeletal and contractile genes, NFκB, and type I interferon-induced genes, as well as cell cycle genes by the target gene knockdowns at both the pathway, marker gene, and regulon analysis levels. Importantly, there is experimental evidence from mouse models that several of these programs in SMCs are causal mediators of atherogenesis. Heterozygous deletion of MYOCD, the transcriptional co-activator of most contractile genes, results in accelerated atherosclerosis with more inflammatory cells in mice ^56^. NFκB signaling in cultured SMCs interferes with MYOCD function and drives proliferation ^57^, and knockout of the gene encoding the required signaling factor IKKβ in SMCs protects against the development of murine atherosclerosis ^58^. Furthermore, we recently found that the depletion of modulated SMCs in regressing atherosclerosis is associated with the loss of NFκB signaling in those cells^8^. In contrast, the role of type I interferon signaling in SMCs and atherosclerosis is less understood ^59^, but the strong regulation observed here, which could not be attributed to artifacts of siRNA transfection, warrants further studies of the potential causal role. Finally, the observed almost uniform downregulation of cell cycle genes and proliferation rates in cell culture assays was striking. SMC proliferation is a key driver of plaque development in atherosclerosis, but it is unlikely that almost all the GWAS genes, including both protective and detrimental ones, regulate proliferation in the same direction *in vivo*. We speculate that our findings can be explained by cultured SMCs being selected for their ability to proliferate - those that proliferate better overgrow others. If the state of the cell is optimized for the fastest possible proliferation, it makes sense that most perturbations to the cell would result in reductions in proliferation. This inherent quality of SMC culture can easily be overlooked when gene knockdowns are examined individually.

A second main finding was that several knockdowns produced different effects in cells subjected to baseline, cholesterol overload, and mechanical stretch conditions. In most cases, the difference was a lack of significant effect in one condition when that effect was present in (an)other condition(s), but some effects were even opposite in different conditions. This is a testament to the importance of the context within which gene function is analyzed, and it touches on a major limitation in the field, further discussed below, that it is not yet known which SMC culture assays are predictive for relevant SMC functions *in vivo*.

Thirdly, we found that commonly used markers for SMC phenotypes and functional readouts like migration and proliferation were independently regulated. This pattern has been reported before ^60^. This contrasts with the clear inverse associations between contractile gene markers (e.g., ACTA2, MYH11) and markers of modulated SMCs (e.g., LUM) and inflammatory signaling in plaque scRNA-seq data ^5,8^, and the fact that healthy contractile SMCs are non-proliferative and non-migrating, whereas activated, non-contractile SMCs in disease expand and move. The likely explanation is that studies in cultured SMCs are confined to examining cellular mechanisms in a semi-modulated, proliferative state, whereas SMCs *in vivo* undergo much more pervasive changes, transitioning from contractile SMCs characterized by low NFκB signaling and minimal synthetic gene expression to non-contractile mesenchymal cell types exhibiting active NFκB signaling and robust synthetic gene expression. These cellular transitioning processes cannot yet be fully modeled in vitro, and none of the knockdowns were consequential enough to transform the overall cell phenotype. This is evidenced by the PCA analysis in **Figure 2D**, which shows that no knockdown shifted gene expression from one cell state cluster to another. The key takeaway is to acknowledge this limitation of SMC culture studies, and not use SMC markers, proliferation, and migration in vitro as if they reported on the full trajectory that SMCs take in atherosclerotic plaques *in vivo*.

The final key finding was that the cholesterol-induced gene response had the characteristics of a pro-atherogenic mechanism. Among the gene knockdowns that regulated cholesterol-induced genes, upregulation was seen after the knockdown of predicted protective genes and downregulation after the knockdown of predicted detrimental genes (**Figure 6**). This type of analysis is only hypothesis-generating, but, importantly, the causal role of cholesterol overloading in SMCs is supported by experimental evidence. Cholesterol overload of cultured SMCs with cyclodextrin-complexed cholesterol, as performed in this study, leads to foam cell formation with cholesteryl ester droplet accumulation ^24^. While this method may not generate cells that fully acquire the transcriptional profile of SMC-derived foam cells in vivo ^26^, it induces gene expression that overlaps with gene expression of modulated SMCs in progressing mouse lesions ^8^. Part of this regulation appears to be mediated by cholesterol overload-induced activation of NFκB signaling ^25^, possibly through eliciting endoplasmic reticulum stress ^61^. It will be important to investigate the upstream and downstream mechanisms of cholesterol overload in SMCs to identify nodes that can be targeted in experimental models to understand causal mechanisms and, potentially, as future targets for therapy. Further explorations of *C1S, LMOD1, FHL1, LRP1, and TIMP2*, identified here as strong regulators of the cholesterol-induced gene response, and the specific gene modules M2 and M21 implicated in the mechanism, can be starting points.

In a general perspective, the approach taken here, combining an in vitro screen with genetic effect predictions, can serve as proof-of-concept for much larger-scale studies in SMCs or other atherosclerosis-relevant cell types to predict the effect of blocking versus stimulating specific mechanisms in atherosclerosis.

### Limitations

There are several important limitations in our study to consider. First, cultured SMCs, even if subjected to cholesterol overload and stretch, do not mimic the full diversity of SMC-derived cell phenotypes in atherosclerosis. However, human aortic smooth muscle cells have been used successfully in previous studies to investigate genetic variations in CAD^48,62^. Second, during the timeframe of this study, we were unable to validate our findings in additional donor samples to evaluate individual variability. Nevertheless, our data provide bona fide mechanistic insights into human SMC biology. Third, the target genes were identified using a relaxed criterion for genome-wide significant association with CAD. Combined with the uncertainty involved in predicting the causal gene at the associated locus, it is possible that some of our target genes are not de facto causal human CAD genes. Fourth, the number of informative GWAS genes on our screen was insufficient for fully exploring the relationships between the effect direction of GWAS genes on human CAD and the mechanisms they regulate in SMCs.

## Conclusions

In summary, our study maps major mechanisms regulated by a set of CAD risk genes with a likely mechanism of action in plaque SMCs. By integrating knowledge of the effect direction of a subset of the analyzed GWAS genes, we identify cholesterol overload of SMCs, as well as specific cholesterol- and stretch-regulated gene modules, as putative pro-atherogenic mechanisms regulated by human genetic predisposition to CAD. The data released with this report include 345 RNA-seq data sets that can be used to explore the mechanisms downstream of 18 SMC-relevant GWAS genes, the effect of cholesterol overload and mechanical stretching on SMC cell state, and interactions between GWAS gene manipulation and SMC state.

## Methods

### Mining of genome-wide association studies (GWAS) datasets

We focused on GWAS from the Cardiovascular Disease Knowledge Portal (CVDKP) ^63^ and Finngen r5 ^64^. Datasets were extracted from CVDKP using Signal Sifter (https://cvd.hugeamp.org/signalsifter.html). We searched for coronary artery disease (CAD) traits, excluding those with lipoproteins (LDL) associations. A ratio was calculated between log10(P value) of CAD/LDL. Hits were included if ratio > 2 and LDL P value > 5e-8. For Finngen datasets, credible sets and top hits were extracted using coronary atherosclerosis (“I9_CORATHER”) as a phenotype of interest, choosing the data tables “Traditional” (with a gene nearest to genomic variant) or “Credible Sets” (where genomic variant is fine mapped to a leading variant and related nearest gene). Genes were removed if already identified by the previous method, or LDL P value ≤ 5e-8 or co-association to lipoprotein traits were previously discovered.

### Retrieving genes assigned to genomic variants

The primary list of genes assigned to the collected genomic variants was compiled as described above. Additionally, each genomic variant was checked using the results of “Variant-to-Gene” (V2G) and “Locus-to-Gene” (L2G) fine-mapping pipelines of the Open Targets Genetics platform ^65^. Evidence in V2G and L2G approaches is based on distance between the variant and the gene transcription start site, and the data from several types of experiments, including molecular phenotype quantitative trait loci (QTL), chromatin interaction, *in silico* functional prediction. A detailed description of these approaches can be found in the documentation of the Open Targets Genetics platform. If any additional gene was identified, it was added to the list.

### Integration of single-cell RNA sequencing studies from public datasets

To estimate the transcriptomic patterns of target genes and gene networks in different cell types, we collected single-cell RNA sequencing (scRNA-seq) data of human atherosclerotic lesions and healthy arteries published previously ^5–7^. Sample information was gathered from the Gene Expression Omnibus database (accession GSE159677, GSE155512 and GSE131778). Raw FASTQ files were downloaded from NIH Sequence Read Archive (SRA) and processed via CellRanger v6.1.2 with 10X Genomics-derived human reference hg38 (2020-A July 7, 2020). Single-cell data filtering, integration, and clustering were done in the Seurat R package ^66^. Cells were filtered as follows: number of UMIs > 500, number of detected genes between 200 and 4500, percent of mitochondrial genes < 10% and hemoglobin genes < 1%, ratio of mitochondrial to ribosomal genes < 0.5 ^67^. Additionally, we discarded the barcodes suspected as doublets using any of the three methods: DoubletFinder ^68^, scDblFinder ^69^, and Scrublet ^70^.

We normalized data using the standard “LogNorm” method in Seurat, and integrated datasets using Harmony ^71^. Cell clusters were identified using a shared nearest neighbor modularity optimization-based algorithm (implemented in Seurat by default) with 2000 most variable genes, 30 principal components, and a resolution of 1. Marker genes of the identified cell clusters were found with Wilcoxon–Mann–Whitney test implemented in Seurat (**Supplementary Table 11**). Automatic cell type annotation was performed using popularVote pipeline (https://tabula-sapiens-portal.ds.czbiohub.org/annotateuserdata) with the Tabula Sapiens “Vasculature” dataset as a reference ^72^. To validate the SMC-specific origin of identified clusters, we integrated human data with a scRNA-seq dataset of SMC-derived cell population from Myh11-CreERT2 x TdTomato (TdT) reporter mice with induced atherosclerosis and high-fat diet as described elsewhere (Carramolino et al, 2024). Cell clusters within the SMC-containing mesenchymal supercluster were further re-annotated based on marker genes (differentially expressed in a specific cell cluster compared to other cell clusters). Amongst the mesenchymal cells, one cluster called “undefined” contained low-quality droplets that passed the filtration step but had dramatically lower numbers of UMIs and detected genes than other cells. This cluster may represent the fragments of mesenchymal cells and, thus, was not considered for further analysis.

### Selection of smooth muscle cell population-specific genes

According to scRNA-seq results, several clusters of SMCs were included in a unified supercluster of “mesenchymal cells.” Since a clear delineation between the spectrum of differentiated and modulated SMCs and other transcriptionally close cell types, including fibroblasts and pericytes, remains challenging, we consider the whole mesenchymal supercluster to determine the SMC specificity of each gene. If a gene had the highest average expression in this supercluster or a gene is expressed in a larger percentage of mesenchymal cells than any other type, or the average expression of a gene in smooth muscle cells was 1.5 standard deviations above the average for all other cell types, we call that gene enriched in mesenchymal supercluster and considered as a candidate gene.

### Spatial transcriptomics

Spatial transcriptomics samples were generated from formalin-fixed and paraffin-embedded (FFPE) human coronary arteries at different stages of the disease, isolated from explanted hearts, as previously described ^20,21^. Briefly, 5µm thickness sections were mounted on Visium slides and processed for spatial transcriptomics according to the 10X Genomics Visium FFPE Version 1 protocol. Samples were deparaffinized, stained with hematoxylin and eosin (H&E), and imaged using VS200 Slide Scanner (Olympus Life Science) before decrosslinking, destaining and overnight probe hybridization with the 10X Visium Human version 1 probe set. The following day, hybridized probes were released from the tissue and ligated to spatially barcoded oligonucleotides on the Visium Gene expression slide. Barcoded ligation products were then amplified and used for the construction of libraries, which consequently were sequenced on a NovaSeq 6000 sequencing platform (Illumina) using a NovaSeq 6000 S2 Reagent Kit v1.5 (Illumina) according to the manufacturer’s instructions. Afterwards, reads were aligned with their corresponding probe sequences, mapped to the Visium spot where each probe was captured, and finally aligned with the original H&E-stained image of the tissue section using the software SpaceRanger version 1.3.0 (10X Genomics). Samples ranged from 568 to 1,746 spots.

### Human aortic smooth muscle cells

We used healthy human aortic smooth muscle cells (haSMCs) from a single donor (ATCC, PCS-100-012; Lot #80323179). SMCs were expanded at 37 °C, 5% CO_2_ in vascular cell basal medium with cell growth kit from ATCC (PCS100 030 and PCS 100 042) from passage P0 to passage P3. During cell expansion, plates were coated with type I collagen from rat tail (Corning, 354236) diluted in 0.1% acetic acid [15 µg/mL] for >20min at room temperature (RT) and washed once with phosphate-buffered saline (PBS). For experiments, SMCs of passage P4 and P5 were cultured in smooth muscle cell growth medium containing 5% fetal calf serum and supplements (Provitro, 211-0601).

### Small interference RNA (siRNA) transfection

The expression of each candidate gene was suppressed by two different small interfering RNAs (siRNAs) per target gene (QIAGEN, 20 μM); the sequences are shown in **Supplementary Table 12**. Following the guidelines provided by the manufacturer, the siRNAs utilized in this study were screened for various sequence motifs known to trigger an interferon or cytokine response. Consequently, siRNAs containing these motifs were excluded from our study. Transfection was performed with Opti-MEM (Thermo Fisher, Cat. No. 31985062) and Lipofectamine RNAiMAX (Invitrogen, Cat. No. 13778150) on two consecutive days according to the manufacturer’s instructions. Control SMCs were transfected with control siRNA (QIAGEN, AllStars Negative S103650318, 20 μM).

### Baseline modulation of human smooth muscle cells

Human SMCs were cultured on type I collagen-coated plastic 12-well plates with growth medium as described above. Cells cultured under these conditions spontaneously modulated their phenotype and thus were used as a “baseline.” When the cells reached 70% confluency, they were transfected with siRNAs for each gene on two consecutive days, as described above. Every RNA sample was a pool of three wells, and we used four technical replicates per condition.

### Cholesterol overload

Human SMCs were cultured on type I collagen-coated plastic 12-well plates with growth medium. At 70% of confluency, cells were either untreated or treated for 72 hours (h) with water-soluble cholesterol (Sigma-Aldrich, C4951, final concentration 10 μg/mL). For initial RNA-seq analysis, we compared expression levels of SMCs treated with cholesterol overload for 72 h versus untreated cells. For target gene characterization, SMCs were treated with cholesterol and transfected in parallel with siRNAs for each gene on two consecutive days while continuing cholesterol treatment for another day to reach 72 h in total. Every RNA sample was a pool of three wells, and we used four technical replicates per condition.

### Mechanical stretch

Human SMCs were seeded onto flexible silicon 6-well plates coated with ProNectin (Bioflex, Cat. No. BF 3001P) and cultured in growth medium until 70% confluency. Cells were then transfected with siRNAs for each gene on two consecutive days, as described above. Then, SMCs were subjected to 10% cyclic stretch with a sinusoidal waveform at a frequency of 1 Hz for 6 h with the Flexcell tension system (Flexcell International Corp., FX-5000 T) or left under static conditions. For initial RNA-seq analysis, we compared expression levels of SMCs subjected to stretch versus untreated cells. For target gene characterization, SMCs were transfected with siRNAs for each gene on two consecutive days. A day later, control and target transfected cells were subjected to stretch. Every RNA sample was a pool of two wells, and we used four technical replicates per condition. Data was evaluated by RNA-seq as described above.

### TNF and IFN treatment

Stimulation with tumor necrosis factor-alpha (TNF) or interferon-alpha (IFN) was performed for 24h when SMCs reached a confluency of ∼80-90% (approximately 48 h after seeding). SMCs were cultured on type I collagen-coated plastic 12-well plates and stimulated with TNF (Abcam, ab259410, 10 ng/mL), IFN (Abcam, ab48750, 100 ng/mL), or left untreated. Every RNA sample was a pool of three wells (12-well plates), and we used four technical replicates per condition.

### Cell proliferation

A total of 50,000 human SMCs were seeded on glass coverslips coated with type I collagen in a 24-well plate. After siRNA transfection, cells were then cultured for an additional 24 h in growth medium containing 5-ethynyl-2’-deoxyuridine (EdU,10 μM). Cells were fixed in 3.7% paraformaldehyde in PBS for 15 min. The EdU-positive cells were labeled according to the manufacturer’s instructions (Sigma-Aldrich, BCK-EDU488), followed by nuclei labeling with DAPI (Thermo Fisher, D1306). Coverslips were mounted on slides and imaged using a Nikon Eclipse Ti2 inverted microscope. The NIS-Elements software was used for image acquisition, and ImageJ ^73^ was used for quantification. Three to four technical replicates were evaluated per gene.

### Scratch assay for migration

Human SMCs were seeded in 24-well plates coated with type I collagen and transfected with siRNAs as described above. Then, the confluent cell monolayer was scratched in the center of the well using a 10 μl pipette tip before washing once with PBS to clear cell debris and floating cells. Cells received fresh media, and the nuclei were labeled with Hoechst 33342 live probe (NucBlue^TM^, Thermo Fisher R37605). Time-lapse images were captured hourly for 24 h using a Nikon Eclipse Ti2 inverted microscope with an integrated chamber ensuring proper cell culture conditions of 37 °C and 5% CO_2_. The migration ability was measured by calculating the percentage of gap closure at different time points compared to time 0 (baseline) using the NIS-Elements software. Three to four technical replicates were evaluated per gene.

### Total mRNA sequencing

RNA was isolated from wells using the NucleoSpin RNA Plus Kit (Macherey-Nagel, 790984) and sequenced by BGI (Copenhagen, Denmark). All RNA samples passed quality control (Agilent 4200 electrophoresis system). Non-stranded cDNA libraries were prepared with polyA-selected mRNA, and 100 bp paired-end sequencing was conducted with at least 20M pair reads per sample using DNBSEQ. RNA sequencing data were processed with the Nextflow-based pipeline “nf-core/rnaseq” (v3.5). Sequenced reads were aligned to the GRCh38 human reference genome using STAR aligner (v2.6.1d), and gene-level quantification was done with Salmon (v1.5.2). The reference genome sequence and gene annotation were downloaded from AWS iGenome using the Ensembl version as a source. Further analysis was performed in R (v4.1.1) with Bioconductor (v3.14) packages. Gene counts were imported using tximport. Gene filtering was performed as previously described, and only genes with ≥10 counts per million (CPM) in at least 3 samples were analysed. DESeq2 ^74^ was used for data normalization, variance stabilizing transformation, principal component analysis, and estimation of differential gene expression between compared groups. Estimated P values (Padj) were adjusted for false discovery rate by Benjamini and Hochberg procedure ^75^. Genes with Padj <0.05 and absolute log2 fold change (LFC) ≥0.5 were considered differentially expressed, or significantly dysregulated. To identify involved pathways, the lists of differentially expressed genes were used for gene-set enrichment analysis using a Web-based gene set analysis toolkit (WebGestalt) ^76^ with categories from GeneOntology ^77^, Reactome ^29^, WikiPathways ^30^, and KEGG PATHWAY ^28^ databases.

### Real time-qPCR

The relative gene expression was measured by real-time qPCR (RT-qPCR) using *HPRT1* as housekeeping gene. Intron-spanning primers amplifying 50<150bp amplicons were designed using the Primer-BLAST design tool from NCBI ^78^. Primer efficiency was tested by plotting the sample quantity of the dilution series to the average ct values of each dilution on a logarithmic scale. The primer sequences are shown in **Supplementary Table 13**. The Maxima SYBR Green qPCR master mix kit [1:2] including ROX [0.18µM] (Thermo Fisher Scientific, K0252) was mixed with forward and reverse primers [268 nM] and cDNA [12.5 ng] in a final volume of 14 µL. On a CFX Opus Biorad machine, the RT-qPCR program was run with an initial denaturation step of 95°C for 10 min followed by 40 cycles of 95°C for 30 sec, 60°C for 1 min, and 72°C for 1 min, and one cycle of 95°C for 1 min, 60°C for 30 sec, and 95°C for 30 sec.

### Regulon enrichment analysis

The lists of unidirectionally dysregulated genes were screened for the enrichment of transcription factor binding motifs using R/Bioconductor package *RcisTarget* ^79^. A set of identified motifs and related transcription factors from TRANSFAC ^80^ database collection was used for further regulon analysis. Regulon activity analysis in the integrated scRNA-seq dataset data of human atherosclerotic lesions and healthy arteries ^5–7^ was performed using the SCENIC pipeline ^79^ with default parameterization and human genome reference (hg38). Area under the curve (AUC) values estimated by the pipeline were averaged for every cell cluster and z-scaled by regulon. Regulons with higher relative activity (positive z-score) in the cluster of interest were considered active.

### Genetic direction of selected CAD-associated genes

We searched for annotated CAD GWAS loci ^81^ and co-localized e/pQTLs collected from several sources (GTEx V8 ^37^, eQTLgen ^38^, UK Biobank ^39^, and deCODE genetics ^40^) for selected target genes. Co-localization analysis was performed using HyPrColoc ^82^ with a posterior probability of alignment threshold of 0.7. For co-localized e/pQTLs, we determined the impact on CAD risk using Mendelian randomization with the e/pQTL as the instrumental variable and the CAD GWAS as the outcome.

### Gene co-expression network inference

We inferred the control gene co-expression network (GCN) using the results of total mRNA sequencing for human SMCs transfected with control siRNA in every cellular assay. Briefly, we employed a gene expression matrix after variance stabilizing transformation. We selected the top 5000 most variable genes for weighted gene co-expression network analysis (Langfelder and Horvath 2008) in BioNERO R package (Almeida-Silva and Venancio 2022). Seven of the 94 control samples were identified as outliers by the standardized connectivity method ^83^ and discarded for further analysis. We used Spearman’s correlation to build a signed network with the power soft threshold 7, achieving scale-free topology R^2^=0.843 (scale-free topology and mean connectivity for different power thresholds are shown in **Supplementary Figure 9**). Correlation analysis between module eigengene (the first principal component of the gene expression matrix of the corresponding module) and cellular state was used to identify gene modules responsive to baseline modulation, cholesterol overload, or mechanical stretch. Noteworthy, we did not adjust RNA sequencing data to correct the difference between experimental batches before GCN inference since they are compromised by study design with a cellular state. Instead, we considered the modules with clear patterns of higher differences between cellular states than within-state variation. Over-representation analysis of genes assigned to different GCN modules in Reactome, KEGG Pathway, WikiPathways, GeneOntology, and Broad Institute Hallmark50 gene categories was performed with WebGestalt, as previously described (**Supplementary Table 10**).

### Cell type deconvolution

We conducted cell-type deconvolution analysis to predict which mesenchymal cell states observed *in vivo* in atherosclerotic lesions can be mirrored by SMCs cultured *in vitro* in our experiments. Transcriptional profiles of mesenchymal cell clusters identified in the integrated scRNA-seq datasets – including contractile, transitional, osteochondrogenic, and pericyte-like SMCs, as well as fibromyocytes and fibroblasts – were utilized as a reference. We estimated the relative proportions of these clusters based on cell counts and generated pseudo-bulk data by summarizing gene counts (using the *AggregateExpression* function in Seurat) in mesenchymal cell supercluster per sample. Then we estimated the relative proportions of mesenchymal cell components in pseudo-bulk samples as well as in every SMC sample from our *in vitro* experiments using the Bisque method ^84^. This method employs scRNA-seq data as a reference to retrieve gene expression profiles and proportions for the given cell type annotation and to learn gene-specific bulk transformations. Non-negative least-squares regression estimates cell proportions from the bulk RNA-seq data. We compared the component proportions between groups of samples using the Wilcoxon–Mann–Whitney test. P values estimated for multiple comparisons were adjusted by the Bonferroni–Holm method.

### Gene signature scoring

We performed gene expression-based scoring of regulation for those gene signatures which, to our knowledge, are related to SMC contractility, adhesion, and cell cycle, as well as with synthesis of components of extracellular matrix and inflammatory signaling. Specific gene sets were collected from pathway databases: KEGG Pathway ^28^, Reactome ^29^, and WikiPathways^30^ (**Supplementary Table 5**). Additionally, we generated sets of top 100 up- and downregulated genes after cholesterol loading, stretch, TNF, or IFN stimulation of SMCs (Padj <0.01, absolute LFC ≥ 1, arranged by LFC) and used them as gene signatures. We also used the gene lists for every module of the SMC-specific GCN as gene signatures (for large modules, only 100 genes with the highest intramodular connectivity were chosen). R/Bioconductor package *singscore* ^85^ was employed for independent sample-wise scoring with DESeq2-produced Wald z-statistic used for gene ranking. Relative z-scores for selected gene signatures were estimated for every comparison of target knockdown versus related control in all studied cellular assays. Thus, the estimated z-scores reflect the change in total transcriptional activity for every gene signature. We assessed the ability of GCN module-specific gene signatures to predict the effect of target genes with identified genetic direction on CAD. Activity score of each GCN module was calculated for every target knockdown and assay, as described above. Random forest model (implemented in the *randomForest* package ^86^) was trained with gene module scores as explanatory variables and the effect of target gene knockdown on CAD as instrumental variable. The embedded measure of node impurity, mean decrease in Gini index, was used to assess how important the contribution of each variable was in the final model prediction.

### Ethics statement

The scRNA-seq datasets were obtained from public studies involving explanted hearts of transplant recipients (for human coronary arteries) and carotid endarterectomies (for human carotid arteries). As stated elsewhere ^5–7^, all patients provided informed consent through forms approved by institutional review boards. The human coronary samples used in spatial transcriptomics were acquired from a commercial supplier (AnaBios Corporation) of tissues donated by organ donors who had consented to donate tissues for research. The analysis involving human coronary arteries (spatial transcriptomics)^20^ was approved by the National Videnskabsetisk Komité and the Regional Ethics Committee in Copenhagen, Denmark. Adhering to local legislation and institutional requirements. Human arterial smooth muscle cells used for the in vitro analysis were obtained from a commercial supplier (American Type Culture Collection, ATCC), where donors signed an informed consent form in accordance with the World Medical Association Declaration of Helsinki on ethical principles for medical research involving human subjects.

## Data availability

All data generated via RNA-seq in siRNA-guided target gene knockdown experiments are deposited at <u>Zenodo</u> and are free to access. As the Danish legislation states, spatial transcriptomics human data can be made available upon reasonable request to the corresponding author and approval from The Danish Data Protection Agency. All other data supporting the findings in this study are included in the main article and associated <u>Supplementary Tables</u>.

## Author contributions

J.A-J., participated in the design of the experiments, performed in vitro experiments, analysis, interpretation of data, and generation of figures, A.M., performed the re-integration of scRNA-seq datasets and RNA-seq bioinformatics analysis, and contributed to the generation of figures, A.L.J., performed in vitro experiments and analysis, P.L.M., performed the initial analysis on GWAS datasets and potential candidate genes, A.K.U., interpreted the data, D.D., contributed to the scRNA-seq analysis and selection of target genes, J.H., performed in vitro experiments and analysis, L.F.J., performed in vitro experiments and analysis, D.S., performed bioinformatics analysis in SCENIC, C.P., and J.M., performed the spatial transcriptomics analysis, G.B., interpreted the data, J.B., interpreted the data, K.H., interpreted the data, L.M.R., provided experimental assistance for in vitro experiments, M.T., performed the genetic analysis of directionality, M.N., contributed in the conceptualization of the project and interpreted the data, Mette.N., conceptualized the project and interpreted the data, J.F.B., conceptualized the project, design the experiments, and interpreted the data. J.A-J., A.M., and J.F.B. drafted the manuscript and figures with important contributions from A.L.J., P.L.M., A.K.U., J.H., L.F.J., D.S., C.P., G.B., J.B., K.H., J.M., M.T., M.N., and Mette.N. All authors read and approved the manuscript.

## Supplemental material

**Supplementary Figure 1:**
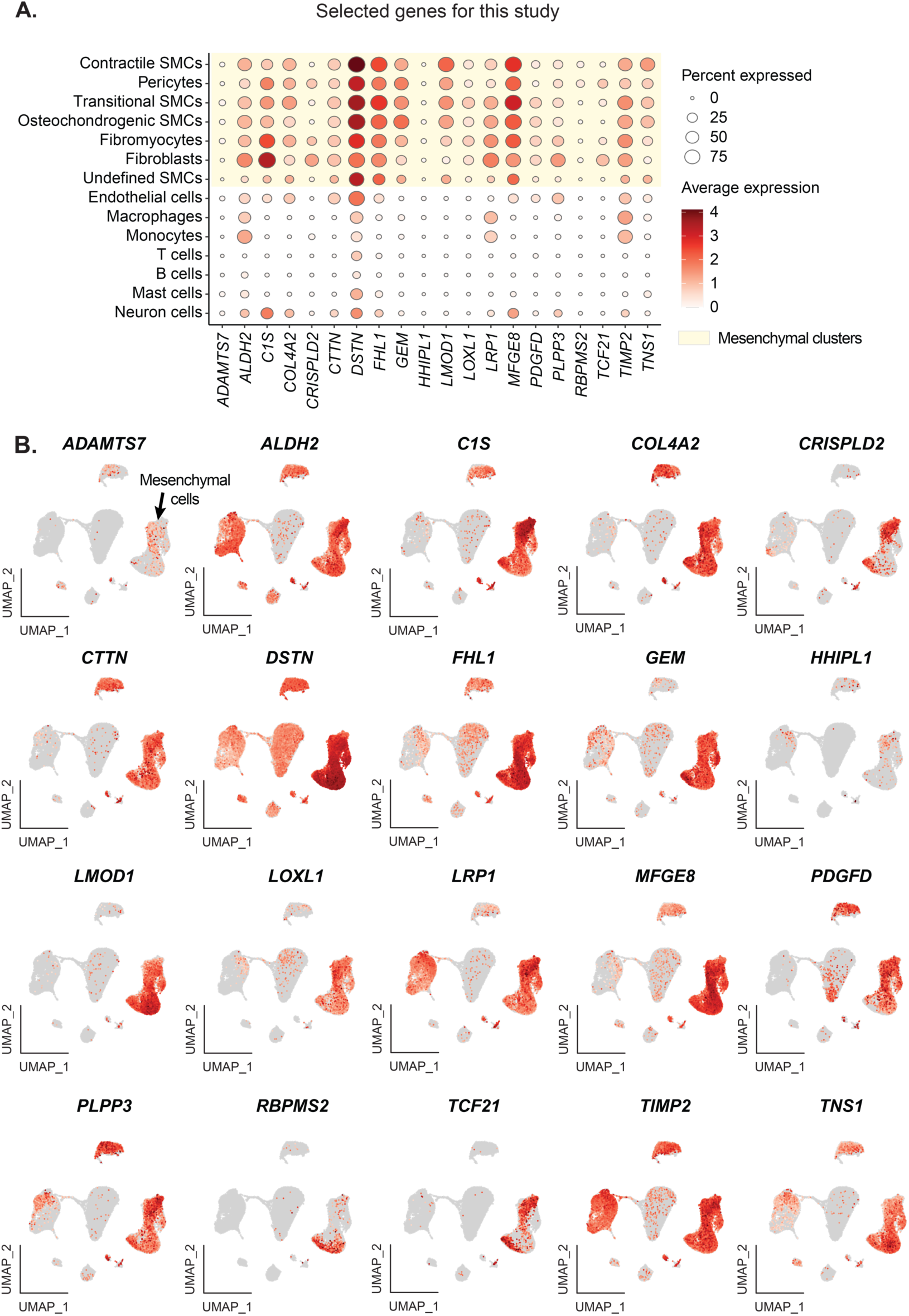
Expression of genes selected for the study in human atherosclerotic plaques. **A.** Dot plot shows the expression of genes enriched in the mesenchymal supercluster (25456 cells) and selected for further study. Normalized and log2-transformed gene expression values are averaged for every cell cluster. **B.** UMAP plots of the selected target genes as analyzed by integrated public scRNA-seq data (50390 cells) on human atherosclerotic lesions. Scale is based on log2-transformed gene expression.

**Supplementary Figure 2:**
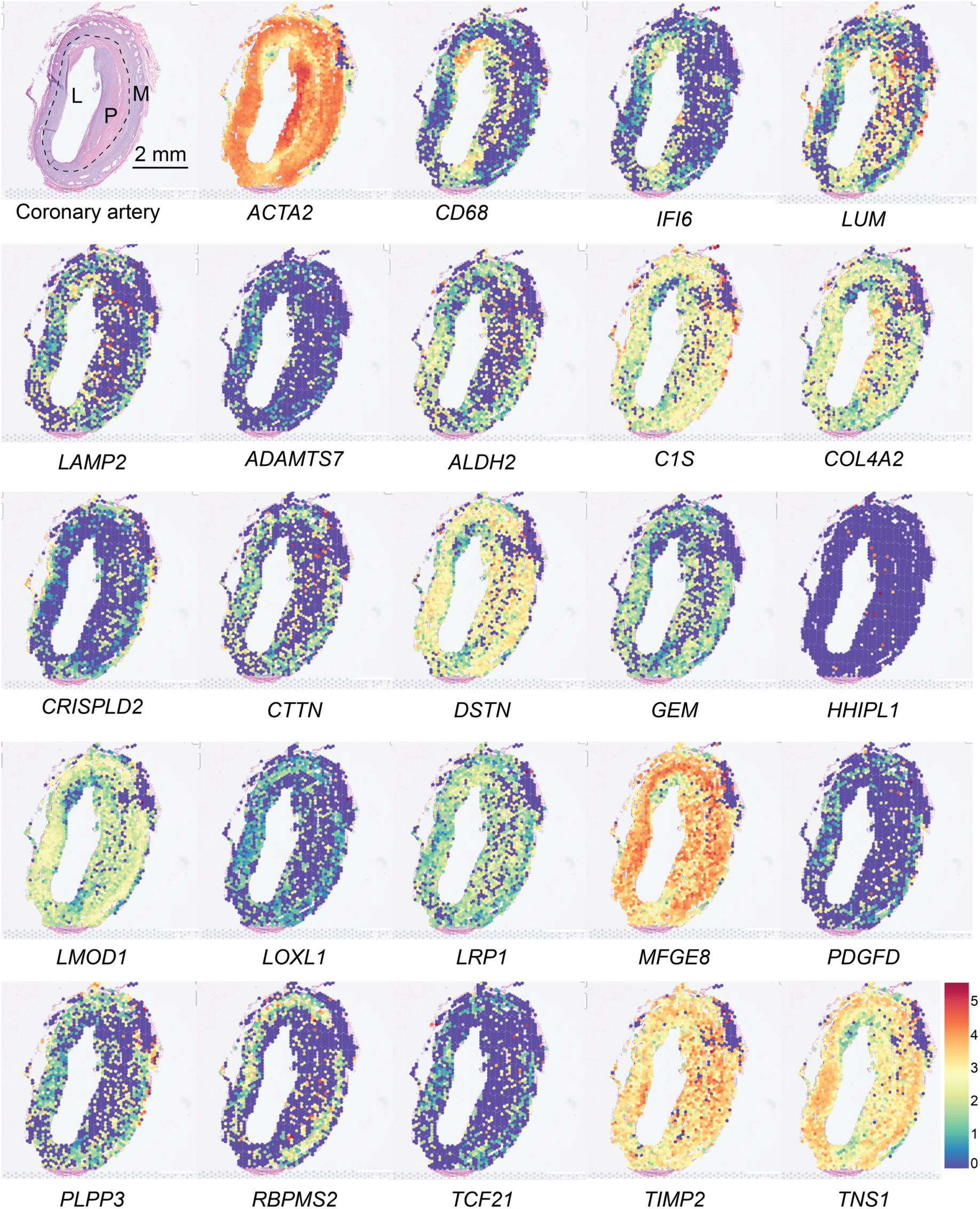
Spatial transcriptomics of selected genes in the study of human coronary arteries with atherosclerotic plaque. Coronary artery plaque is shown in H&E staining *top left* M (media), P (plaque), and L (lumen). Selected markers are shown to identify cell populations (smooth muscle cells; *ACTA2*, macrophages; *CD68*, inflammatory; *IFI6*, modulated smooth muscle cells; *LUM*, and macrophage-like *LAMP2*). Barcoded spots represent a mixture between 5 and 10 cells. Shown is a representative section that contains 1746 spots. Expression intensity is designated by the scale color.

**Supplementary Figure 3:**
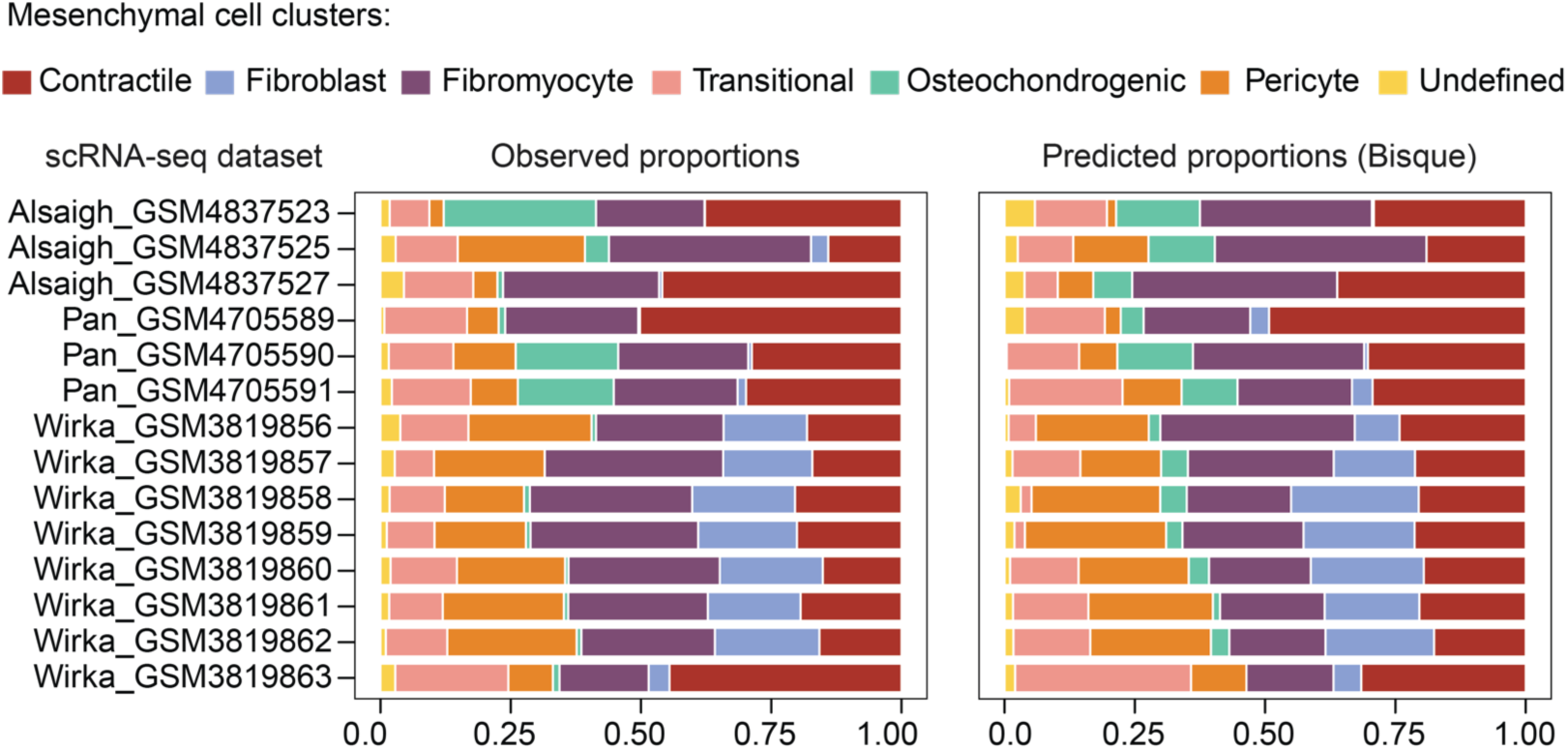
Mesenchymal cell type deconvolution. Proportions of mesenchymal cell clusters observed in scRNA-seq data of human atherosclerotic plaques (left panel) and proportions predicted by cell type deconvolution of pseudo-bulk RNA sequencing data based on by-sample aggregation of gene counts across mesenchymal clusters of the same scRNA-seq dataset (right panel).

**Supplementary Figure 4:**
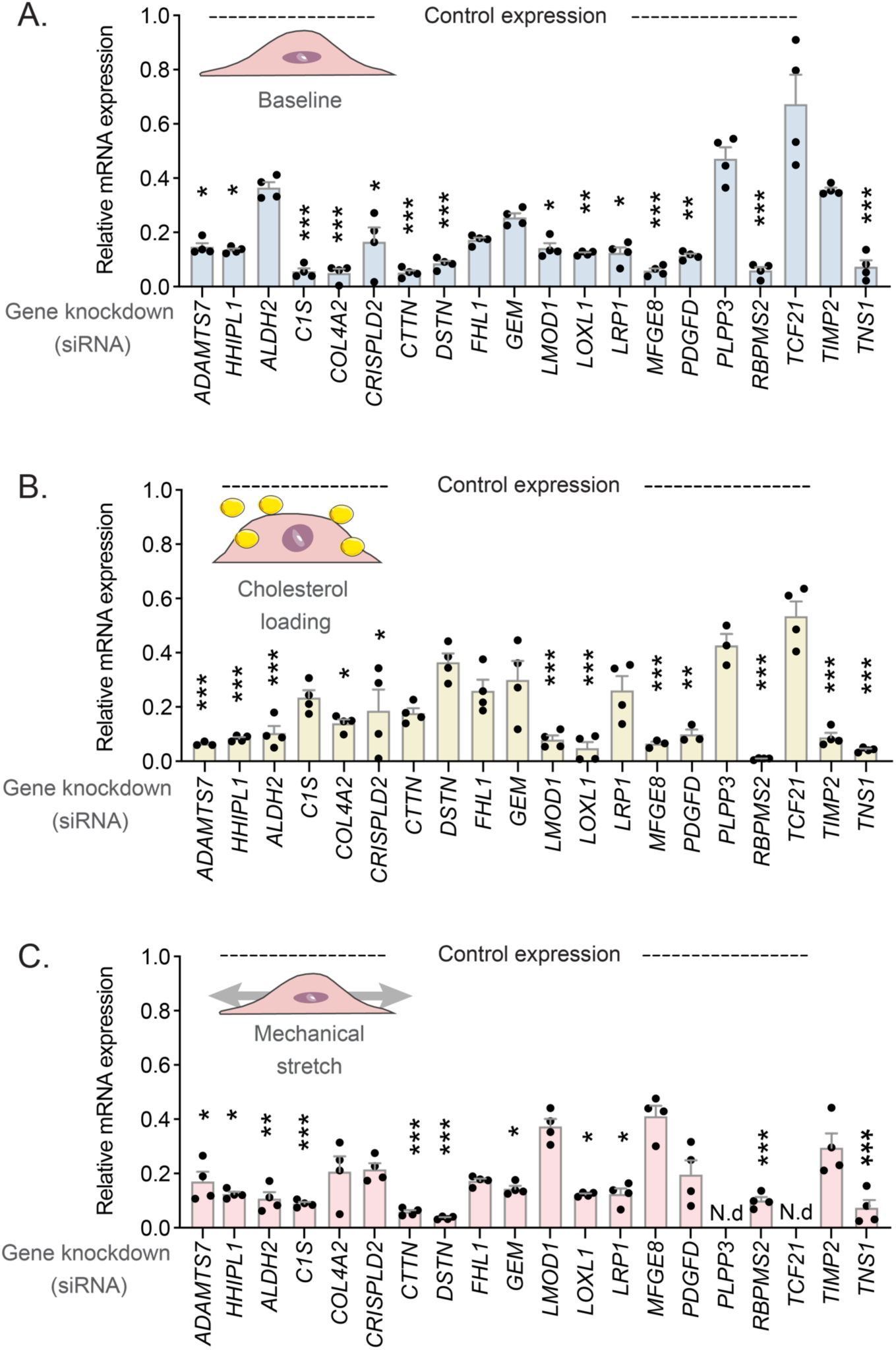
Validation of gene knockdown by real-time qPCR. The knockdown efficiency of target genes in SMCs was evaluated by real-time qPCR under baseline (**A**), cholesterol loading (**B**), and mechanical stretch (**C**) conditions. Gene expression was normalized to the *HPRT1* housekeeping gene (control expression). N.d (gene expression was not determined). **(A-C)** Four technical replicates were tested for each gene. * Adjusted P value < 0.05, ** P value < 0.01; and *** P value < 0.001 compared to control. Comparisons were analyzed by the Kruskal-Wallis test followed by Dunn’s posthoc test with a control of false discovery rate by Benjamini and Hochberg method.

**Supplementary Figure 5.**
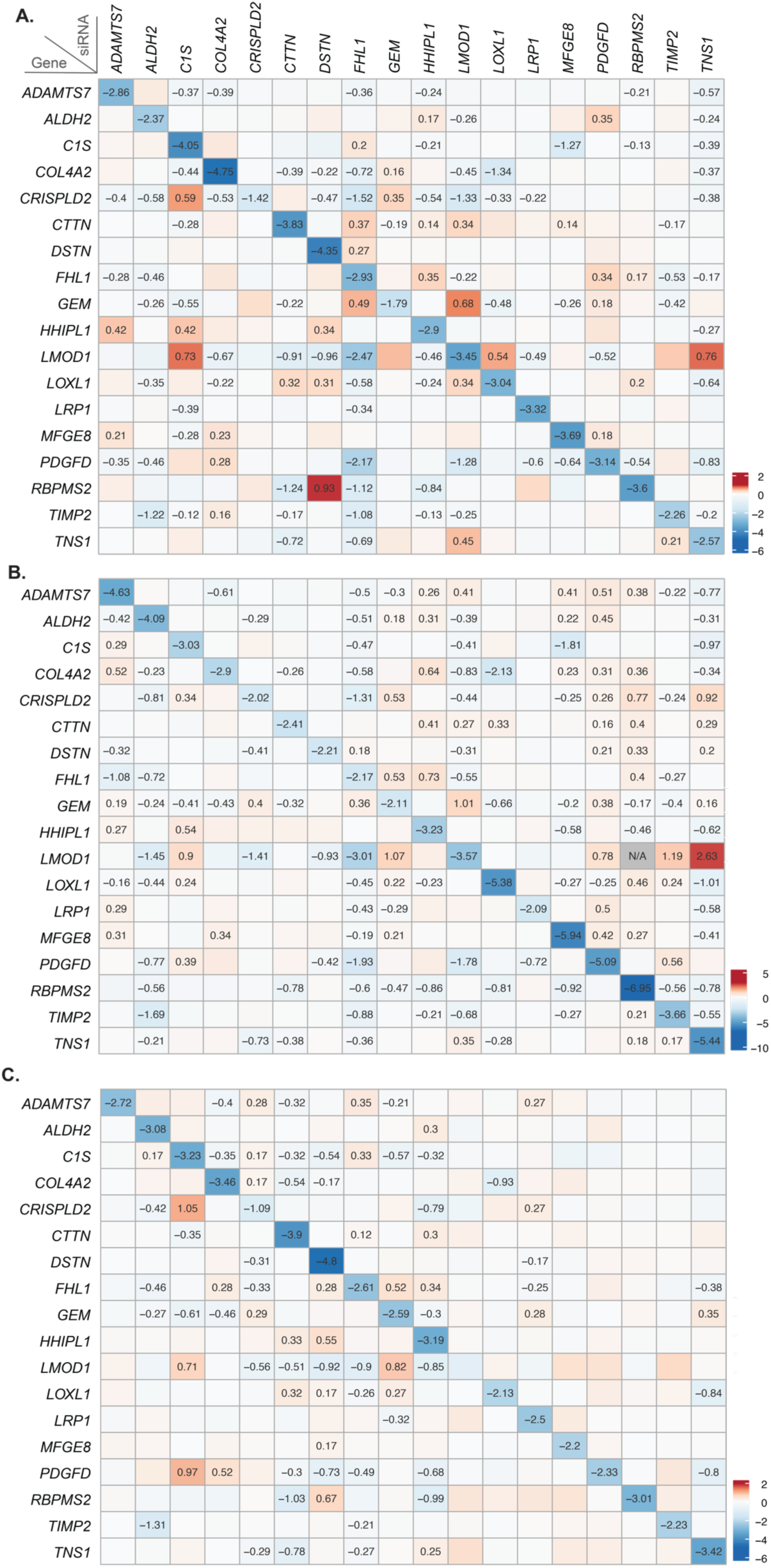
Knockdown efficiency of target genes by RNA-seq. The knockdown efficiency of target genes was evaluated by RNA-seq analysis after baseline (**A**), cholesterol overloading (**B**), and mechanical stretch (**C**) conditions (3 or 4 technical replicates were used for every group of comparison). Genes that are knocked down with siRNAs are located on the X-axis. The measured gene expression is on the Y-axis. Scale is based on log2 transformed fold-change values, and numbers are shown for only significant results with an adjusted P value < 0.05. N/A, the values are not available.

**Supplementary Figure 6:**
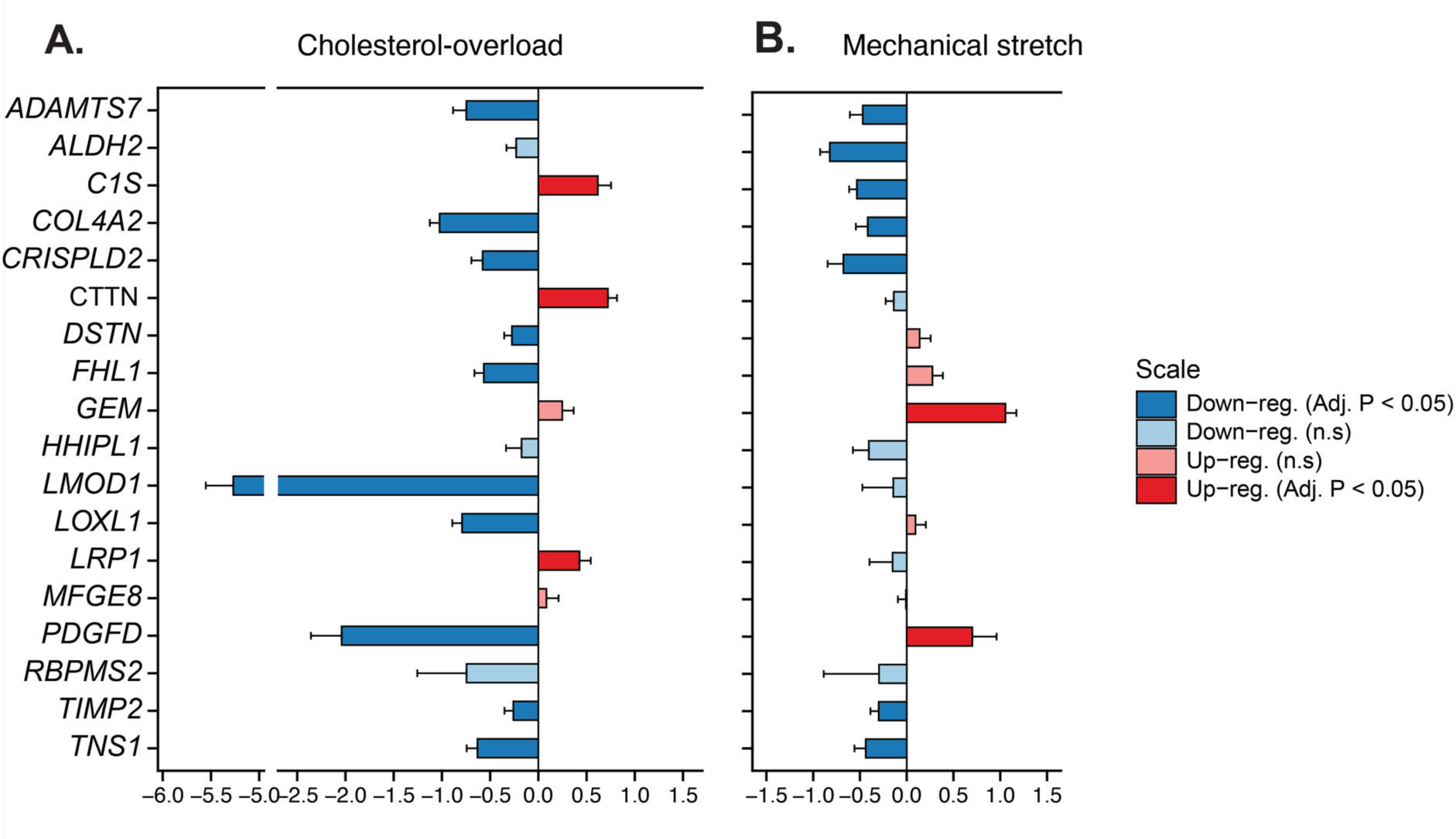
Regulation of target genes in the different cellular assays. Differences in target gene expression in cholesterol overload (**A**) and mechanical stretch (**B**) assays compared to baseline (3 technical replicates in each group). Bars represent log2-transformed fold change (LFC) of average gene expression, error bars are estimated standard error values of LFC. The color scale indicates the direction of fold change (up- and down-regulated genes), significant results (P value adjusted for false discovery rate < 0.05) are highlighted with a saturated color. n.s indicates not significant values.

**Supplementary Figure 7:**
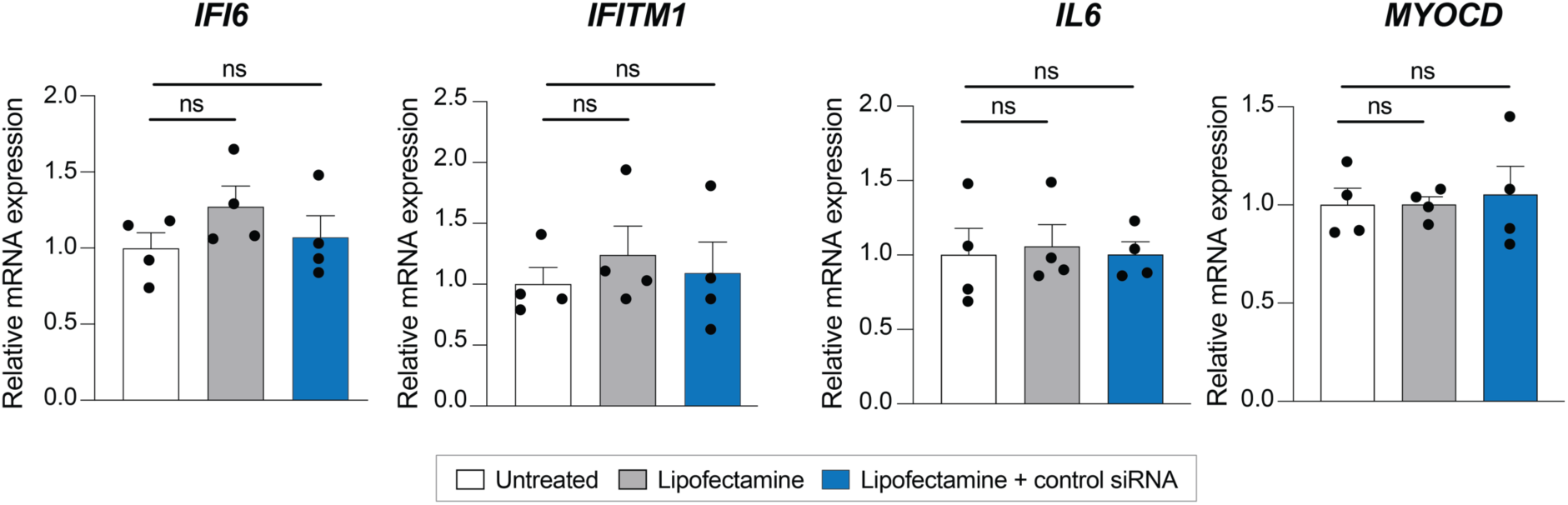
Effect of lipofectamine and control siRNA *per se* in knockdown experiments. Shown is the expression of interferon-regulated genes (*IFI6, IFITM1*), TNF-regulated (*IL6*), or a prototypical contractile gene marker (*MYOCD*) in SMCs treated with lipofectamine, control siRNA or left untreated. Gene expression was normalized to the housekeeping gene *HPRT1*. Four technical replicates were tested for each gene. Statistical significance was tested by one-way ANOVA. Non-significant results (P > 0.05) are shown as ns.

**Supplementary Figure 8:**
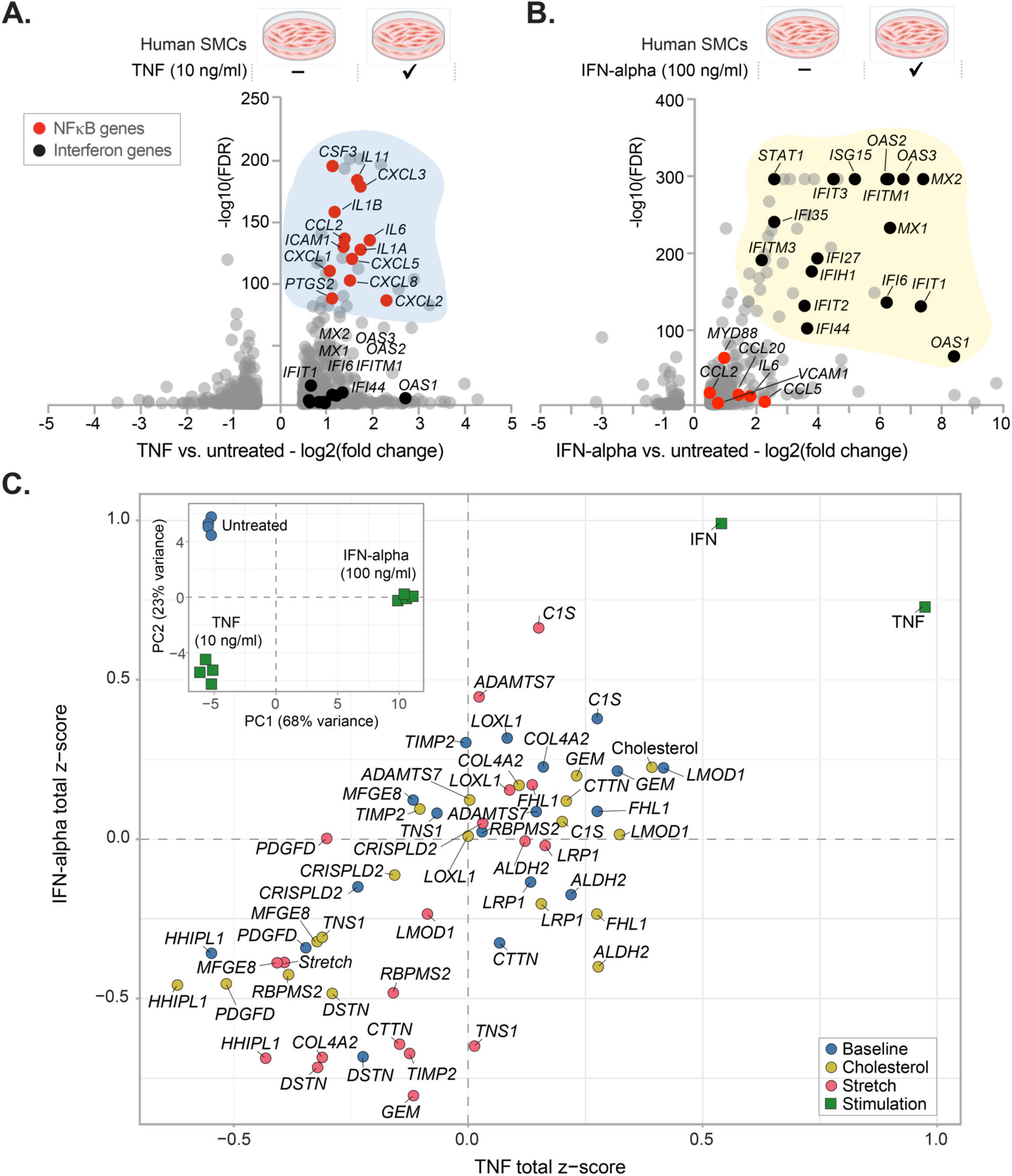
NFκB and type I interferon gene signatures in human SMCs. Transcriptomic changes in human SMCs treated with **(A)** tumor necrosis factor (TNF, 10 ng/ml) and with **(B)** interferon alpha (IFN-alpha, 100 ng/ml) for 24h compared to untreated cells (4 technical replicates in compared groups). Shown in A and B are DEGs with a P value < 0.05 and absolute log2 fold-change ≥ 0.5. **C.** To understand how NFκB and type I interferon gene networks are regulated by selected CAD genes, we scored the alterations in the expression of TNF and IFN-alpha gene signatures in every target gene knockdown and cellular assay (baseline, cholesterol overload, or stretch). The scores of the TNF and IFN-alpha stimulated cells compared to unstimulated control are added for reference (green square symbols). The inserted plot in **C** shows the principal component analysis (PCA) of cells stimulated with TNF, IFN-alpha, or left untreated. Some elements in A and B were created with BioRender.com

**Supplementary Figure 9:**
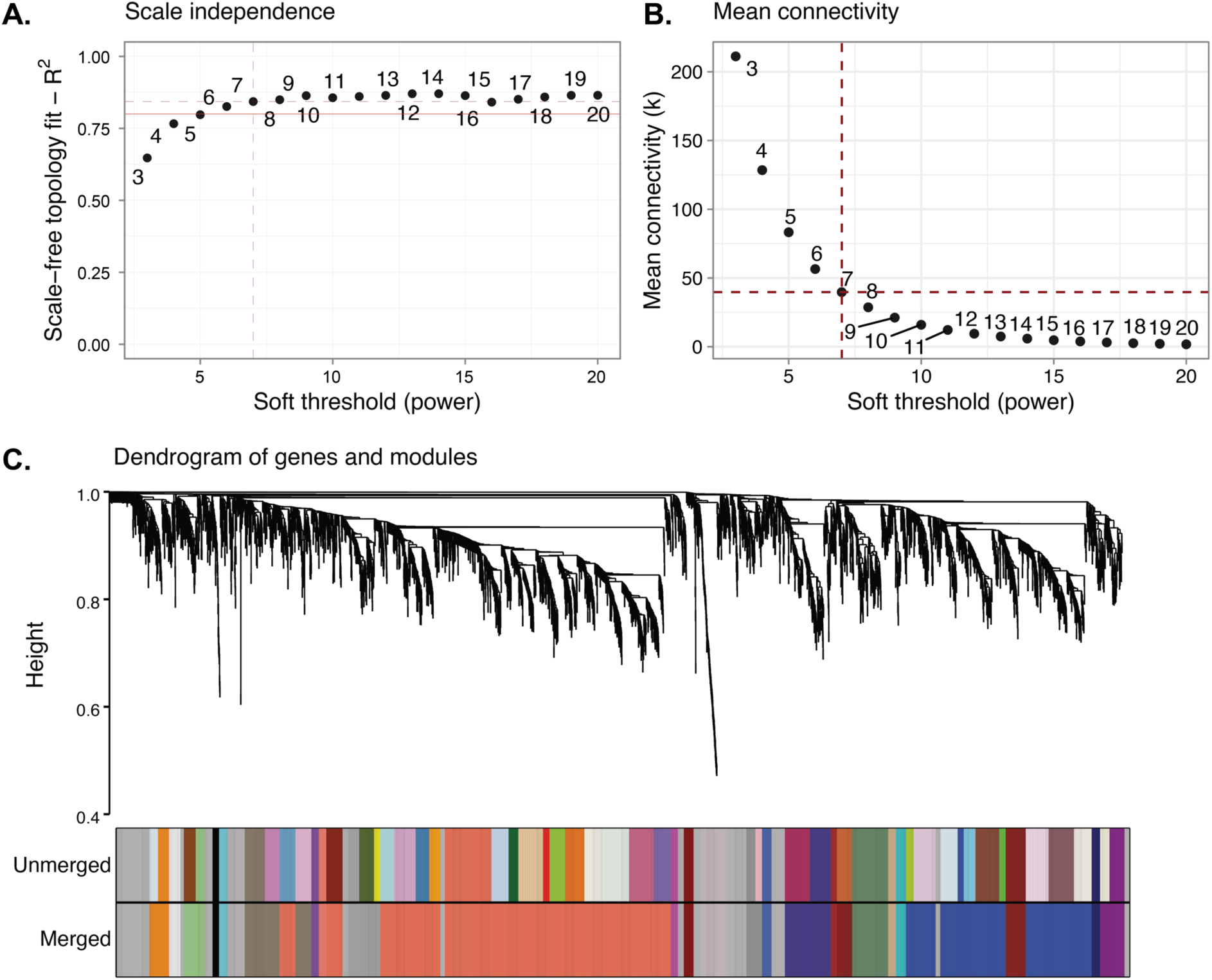
Gene co-expression network analysis in control human aortic smooth muscle cells. **A** and **B** show the soft threshold power analysis results to ensure an optimal scale-free model fit index and mean connectivity, respectively, for weighted correlation network analysis of 94 samples of control siRNA-treated cells in our study. The red solid line demarcates a lower cutoff for the fit index (R^2=0.8). Dashed lines highlight the selected soft threshold**. C.** Dendrogram plot of 5000 genes in a co-expression network clustered based on a dissimilarity measure and their assignment to different modules (shown as different colors).

**Supplementary Figure 10.**
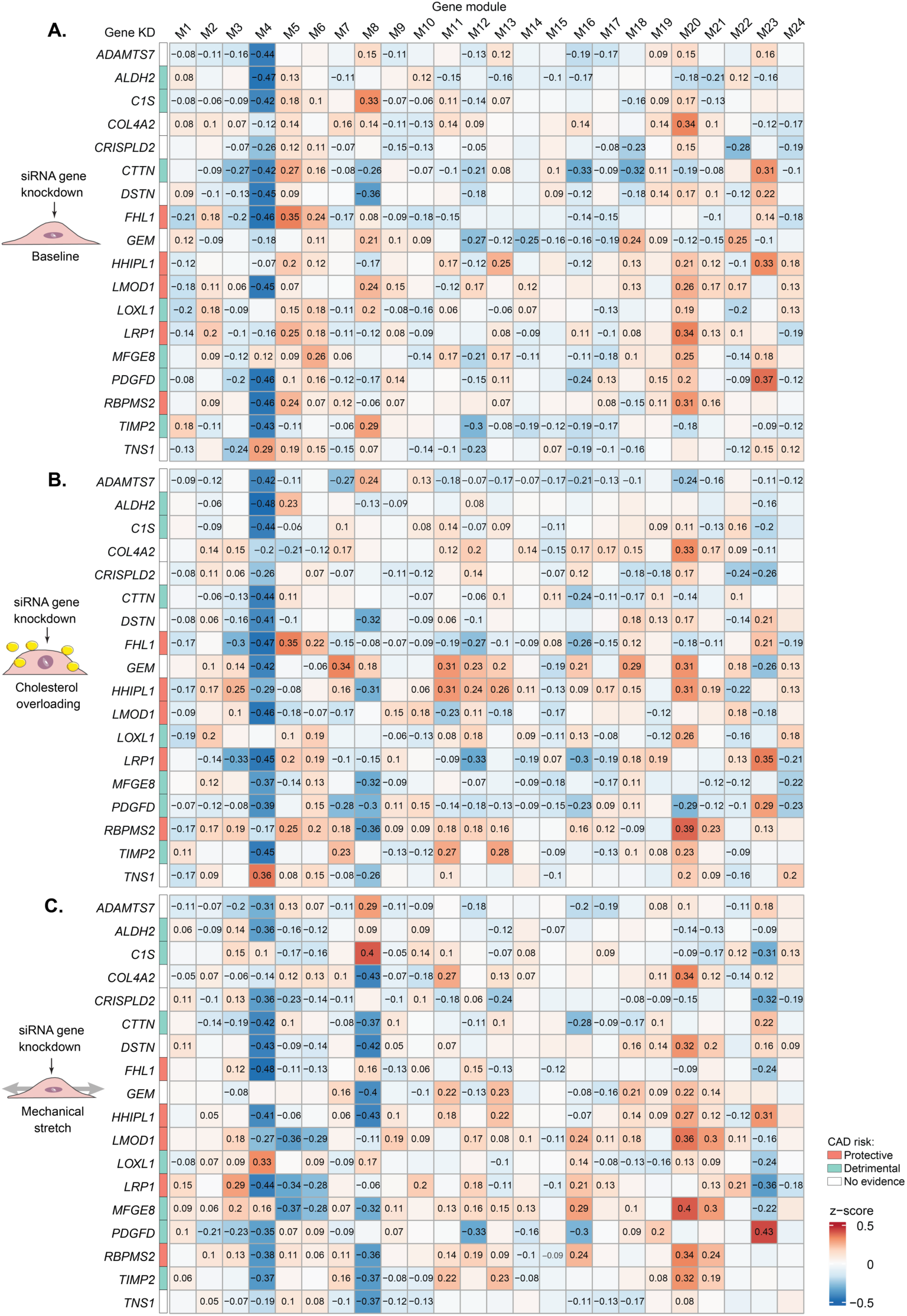
Gene modules co-expressed in human SMCs and disturbed after target gene knockdown. Expression-based scoring of gene signatures for SMC-specific modules of co-expressed genes in the different cellular assays: **A.** baseline, **B.** cholesterol overload, and **C.** mechanical stretch. For gene modules 1 to 10, only the top 100 genes with the highest intramodular connectivity are considered for this figure. The color scale represents a z-score of deviation in the transcriptional activity of the relevant module’s gene signature in target knockdown compared with a control group (3 or 4 technical replicates in each group). Only statistically significant z-score values are shown. Genetic direction is shown for genes associated with a decreased risk for coronary artery disease (CAD, in salmon color), increased (emerald color), or unknown (white color). Gene knockdown (KD).

## Supplementary Tables

Supplementary Table 1 Target list of CAD genes identified after LDL-cholesterol exclusion.

Supplementary Table 2 Genes up-regulated in every identified cell cluster compared to other cells of integrated plaque scRNA-seq data

Supplementary Table 3 List of target genes with methods used for selection

Supplementary Table 4 Gene-set enrichment analysis of differentially expressed genes

Supplementary Table 5 Gene signatures used to score relevant pathways in smooth muscle cells

Supplementary Table 6 TNF- and IFN-regulated genes

Supplementary Table 7 Regulons up- and down-regulated per condition

Supplementary Table 8 Target gene directionality

Supplementary Table 9 Modules of gene correlation network

Supplementary Table 10 Enrichment analysis of genes with the highest intramodular connectivity (up to 100 genes) in the modules of gene co-expression network

Supplementary Table 11 Gene markers of cell clusters identified in integrated scRNA-seq data of human atherosclerotic lesions

Supplementary Table 12 Sequences of small interfering RNA (siRNA) used for target gene knockdown

Supplementary Table 13 Primer sequences used for RT-qPCR

